# Identification and characterization of inflammatory LILR and fibrotic SPP1 macrophages in chronic lung allograft dysfunction

**DOI:** 10.64898/2026.09.04.749490

**Authors:** Allen Duong, Sajad Moshkelgosha, Aaron Wong, Ankita Burman, Ke Fan Bei, Rayoun Ramendra, Tsukasa Ishiwata, Boris Hinz, Sonya MacParland, Mingyao Liu, Stephen Juvet, Tereza Martinu

## Abstract

Chronic lung allograft dysfunction (CLAD) is a major cause of death after lung transplantation and manifests principally as bronchiolitis obliterans syndrome (BOS) or restrictive allograft syndrome (RAS). Although pulmonary macrophages are key regulators of lung injury and repair, their contributions to CLAD pathogenesis remain under-examined. We performed single-cell RNA sequencing-based transcriptomic analysis of CLAD (7 BOS, 7 RAS) and control lung tissue (n = 6), complemented by comparative and functional analyses. Two distinct macrophage populations were identified in CLAD: inflammatory macrophages expressing leukocyte immunoglobulin-like receptors (LILR), and fibrotic macrophages characterized by expression of osteopontin (SPP1). LILR macrophages were present in both BOS and RAS, whereas SPP1 macrophages were selectively enriched in RAS. Both populations were identified in idiopathic pulmonary fibrosis and cross-tissue comparisons. Surface marker-based sorting strategies were developed to isolate both populations. Functional studies demonstrated that LILR macrophages exhibited enhanced phagocytic activity, promoted T cell chemotaxis and secreted inflammatory mediators, whereas SPP1 macrophages produced soluble factors that drove fibroblast activation and contraction. These findings identify distinct inflammatory and fibrotic macrophage programs associated with CLAD, link RAS to conserved fibrotic macrophage states observed in pulmonary fibrosis and highlight macrophage populations as potential therapeutic targets for inflammatory and fibrotic allograft injury.

## INTRODUCTION

Chronic lung allograft dysfunction (CLAD) is the main cause of death for lung transplant recipients, limiting median post-transplant survival to approximately six years (1). CLAD is characterized by a progressive and irreversible decline in lung function and encompasses two major clinical phenotypes with distinct physiological and pathological features: bronchiolitis obliterans syndrome (BOS) and restrictive allograft syndrome (RAS) (1, 2). BOS manifests with obstructive physiology and is pathologically characterized by obliterative bronchiolitis with small airway inflammation and fibrosis. In contrast, RAS presents with restrictive physiology, is associated with pleuroparenchymal fibroelastosis, and is linked with significantly worse clinical outcomes. Despite clear clinical and pathological distinctions between BOS and RAS, the mechanisms driving CLAD pathogenesis and the histopathological divergence of these syndromes remains poorly understood. Additionally, a mix of BOS and RAS pathological features often exist within CLAD allografts, suggesting a spectrum of fibrotic severity may be at play.

Pulmonary macrophages have gained increasing recognition as key regulators of lung injury and repair in lung diseases and pre- and post-transplantation (3–7). These cells exhibit remarkable plasticity and assume functions that are highly dependent on their local microenvironment (6, 8). Under steady-state conditions, macrophages contribute to host defense and immune homeostasis (9), whereas during inflammatory states they adopt enhanced phagocytic activity (10, 11) and secrete pro-inflammatory cytokines that shape alloimmune T cell responses (12, 13). In addition, macrophages play a direct role in tissue fibrosis by producing pro-fibrotic mediators, including transforming growth factor-beta (TGFβ) and modulating fibroblast activation and function (14–18). Given that inflammation and fibrosis are central features of CLAD pathogenesis (19), a detailed understanding of macrophage biology in this context is critically needed.

In lung transplantation, resident alveolar macrophages are key contributors to ischemia-reperfusion injury (20, 21); however, their role in the development of CLAD is less well defined. Using single-cell RNA-sequencing (scRNAseq), we previously identified transcriptionally and phenotypically distinct AMΦs in bronchoalveolar lavage (BAL) samples associated with acute lung allograft dysfunction and in a small cohort of BOS-type CLAD (22). Nevertheless, BAL sampling is restricted to the alveolar compartment and may not adequately assess interstitial macrophages (IMΦ) in the parenchyma. Therefore, comprehensive assessment of MΦ heterogeneity and function in CLAD requires direct analysis of lung tissue.

In this study, we performed scRNAseq on single-cell suspensions generated from end-stage CLAD lung tissue from patients with BOS or RAS and compared these samples with non-CLAD control lungs. We identified two distinct macrophage populations: a pro-inflammatory subset expressing leukocyte immunoglobulin-like receptors (*LILR*), here named LILR macrophages and a pro-fibrotic subset characterized by osteopontin (secreted phosphoprotein 1; *SPP1)* expression, called SPP1 macrophages. While LILR macrophages are preferentially associated with BOS, SPP1 macrophages characterize RAS. Using complementary *in vitro* functional assays, we demonstrate that these populations differentially mediate T cell chemotaxis, modulate fibroblast activity and secrete inflammatory and fibrotic cytokines and chemokines. Together these findings suggest that distinct macrophage programs play divergent and pathogenic roles in CLAD pathogenesis.

## RESULTS

### Cellular composition changes in CLAD

Fourteen CLAD lung samples comprising BOS (n = 7) and RAS (n = 7) phenotypes were included for scRNAseq (Table 1). One patient with undefined CLAD phenotype and no RAS-like opacities on chest imaging was classified as BOS. Four patients with mixed CLAD phenotype and RAS-like fibrotic opacities on chest imaging were included in the RAS group, given the presence of parenchymal fibrosis and similarity in outcomes (23). Recipient demographics, cytomegalovirus serostatus matching, immunosuppression and donor-specific antibody prevalence were similar between groups, whereas pulmonary fibrosis as the native disease, and shorter time to death or retransplant were more common in RAS. Active infection at tissue procurement were observed in both cohorts.

**Table 1.** scRNAseq Cohort Patient Characteristics.

|  | BOS | RAS | P value |
| --- | --- | --- | --- |
|  | n = 7 | n = 7 |  |
| CLAD phenotype | BOS = 6 | RAS = 3 |  |
|  | Undefined = 1 | Mixed = 4 |  |
| Recipient age at transplant (mean) | 34.3 ± 14.1 | 39.0 ± 15.8 | 0.6422 |
| Male n (%) | 1 (14.3%) | 3 (42.9%) | 0.5594 |
| Tissue obtained through |  |  | 0.4615 |
| Re-transplant | 5 (71.4%) | 7 (100%) |  |
| Autopsy | 2 (28.6%) | 0 (0%) |  |
| Time from transplant to CLAD (days) | 1137 ± 897 | 617 ± 296 | 0.3456 |
| Time from transplant to death/re-transplant (days) | 2268 ± 1063 | 1480 ± 642 | 0.1282 |
| Time from CLAD to death/re-transplant (days) | 1131 ± 853 | 898 ± 743 | 0.6282 |
| Native lung disease |  |  | 0.1406 |
| Pulmonary Fibrosis | 0 (0%) | 3 (42.9%) |  |
| COPD/Emphysema | 1 (14.3%) | 2 (28.6%) |  |
| Cystic Fibrosis | 2 (28.6%) | 1 (14.3%) |  |
| Other* | 4 (57.1%) | 1 (14.3%) |  |
| Sex matching (R/D) (n) † |  |  | 0.5850 |
| M/M | 1 | 3 |  |
| F/F | 4 | 3 |  |
| M/F | 0 | 0 |  |
| F/M | 1 | 1 |  |
| CMV serostatus matching (n) |  |  | 0.5850 |
| R-/D+ | 3 (42.9%) | 2 (28.6%) |  |
| R+/D+, R+/D- | 2 (28.6%) | 4 (57.1%) |  |
| R-/R- | 2 (28.6%) | 1 (14.3%) |  |
| Infection at time of tissue procurement‡ | 2 (28.6%) | 3 (42.9%) |  |
| DSA status |  |  |  |
| At time of tissue procurement | 2 (28.6%) | 1 (14.3%) | 0.5148 |
| Historic | 5 (71.4%) | 5 (71.4%) | >0.9999 |
| Immunosuppressant at time of tissue procurement<sup>§</sup> |  |  |  |
| Calcineurin inhibitor (n) |  |  | 0.9056 |
| Tacrolimus | 6 | 5 |  |
| Cyclosporin | 1 | 1 |  |
| Cell cycle inhibitor (n) |  |  | 0.3352 |
| Azathioprine | 1 | 0 |  |
| Mycophenolic acid/mycophenolate mofetil | 6 | 6 |  |
| Dose of prednisone, median (range) [mg/day] | 11.43 (5-20) | 12.92 (10-17.5) | 0.7176 |
| Azithromycin use at time of tissue procurement<sup>‡</sup> |  |  |  |
| Yes | 7 | 6 |  |
| No | 0 | 0 |  |
| Acute Rejection |  |  |  |
| A score, median (range) | 0.67 (0 – 3) | 0.22 (0 – 0.6) | 0.5541 |
| B score, median (range) | 0.04 (0 – 0.25) | 0.07 (0 – 0.33) | 0.7273 |

\*Other primary lung diseases included: Eisenmenger's syndrome (BOS, n = 1), lymphangioleiomyomatosis (BOS, n = 1), sarcoidosis (BOS, n = 1), pulmonary arterial hypertension (BOS, n = 1; RAS, n = 1).
†Missing data for 1 donor
‡ Presence of pathogen and initiation of targeted treatment at time of explant and within the preceding 2 weeks, or colonization with pathogen at time of explant
§Missing data for 1 recipient
A score, grade of acute cellular rejection based on perivascular and interstitial inflammation according to the International Society for Heart and Lung Transplantation (ISHLT) grading system. B score, grade of airway-centered inflammation (lymphocytic bronchiolitis) according to the ISHLT grading system.
Abbreviations: CLAD, chronic lung allograft dysfunction; COPD, chronic obstructive pulmonary disease; D, donor; DSA, donor-specific antibodies; R, recipient.

Explanted CLAD lung tissue was dissociated into single-cell suspensions and subjected to scRNAseq. Sequencing quality-control metrics for the 14 CLAD samples, including number of cells recovered, mean reads per cell, median genes and UMI counts per cell, sequencing saturation and fraction of reads in cells are summarized in Supplemental Table 1. The CLAD samples were compared with control samples (n = 6) obtained from donor lungs in a previous study (21) to identify disease-associated cellular states (Figure 1A). Integrated clustering identified major immune and parenchymal cell populations in all groups (Figure 1B, Figure S1A-B) with minimal batch effects across samples (Figure S2A). Cell-type annotations were supported by expression of established marker genes, with median and mean expression across annotated cell populations summarized in Supplemental Table 2.

**Figure 1:**
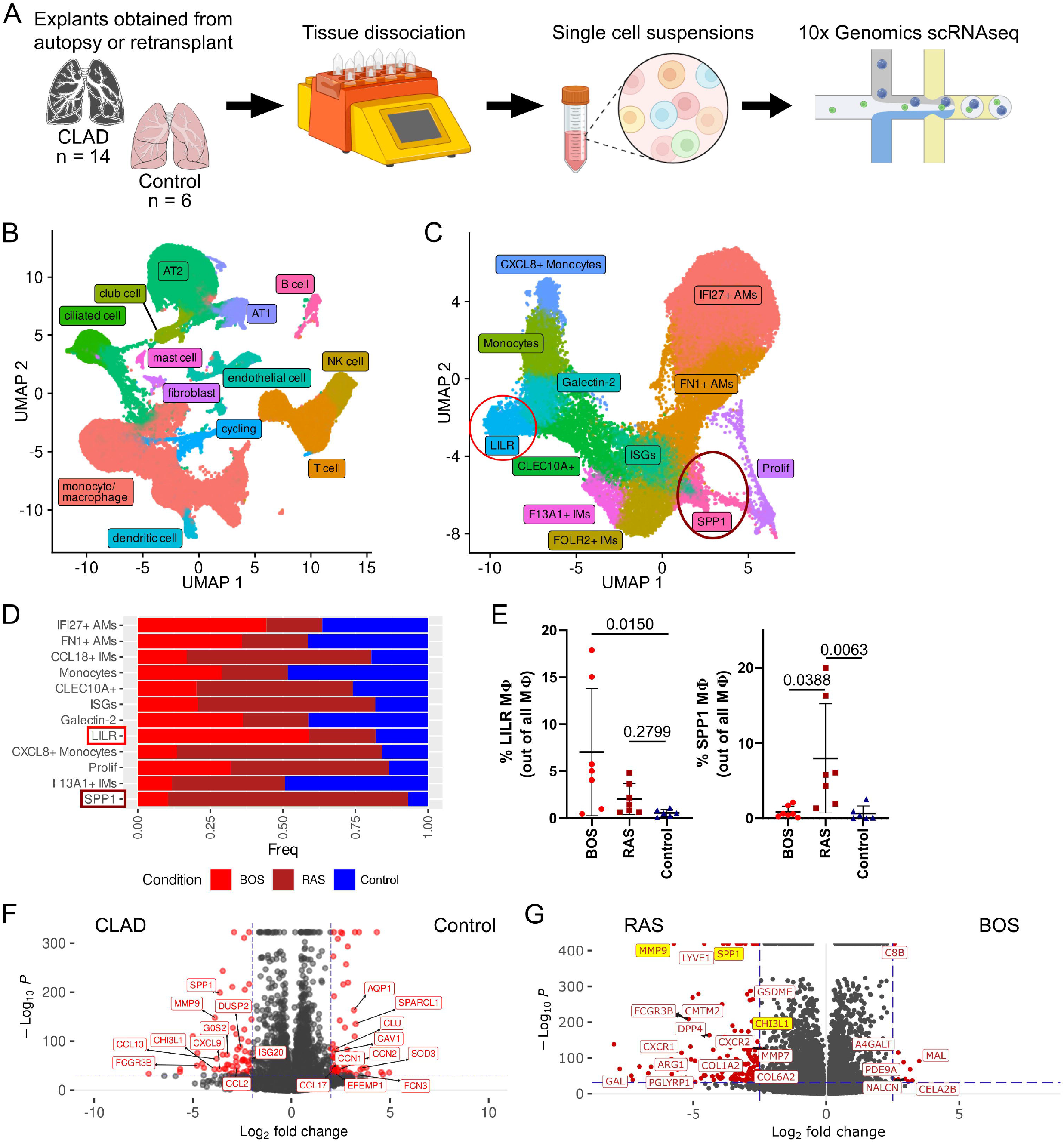
Single-cell RNA sequencing analysis of CLAD lung tissue identifies LILR and SPP1 macrophages enriched in BOS and RAS. A) Single-cell RNA sequencing was performed on single cell suspensions from 14 CLAD lungs (BOS = 7, RAS = 7) and 6 control lungs. B) Single cells from all lungs were integrated, clustered and visualized as an annotated Uniform Manifold Approximation Projection (UMAP) plot. C) Cells defined as monocyte/macrophages were subsetted and reclustered, generating a UMAP plot of 12 annotated macrophage subsets. D) Stacked bar plot showing the proportional distribution of each macrophage subset across BOS, RAS, and Control lungs. E) Proportional representation of LILR and SPP1 macrophages across individual lung samples. Group comparisons were performed using a Kruskal-Wallis test, with post hoc multiple comparisons-adjusted P values indicated in the figure. F and G) Volcano plots showing differential gene expression in macrophages identified by model-based analysis of single-cell transcriptomics (MAST) analysis between F) CLAD versus Control and G) RAS versus BOS. Genes meeting significance thresholds (adjusted *P* value < 0.05, log₂ fold change > 2.5) were highlighted in red. *XIST* was excluded from the volcano plot because its differential expression reflected sex differences between CLAD and control samples rather than disease-associated transcriptional changes.

Cell proportion analysis demonstrated significant enrichment of T cells and a concomitant reduction in type I and type II alveolar epithelial cells in both BOS and RAS compared with controls (Figure S2B-D). Monocytes/macrophages constituted the dominant immune population in CLAD lungs, and were significantly increased in BOS relative to controls (Figure S2D), prompting focused downstream analysis of this compartment.

### Macrophage heterogeneity in CLAD lungs

Cells annotated as “Monocytes/macrophages” were subsetted from the initial integrated dataset and reclustered to assess intrapopulation heterogeneity. At low-resolution reclustering (0.1), four major macrophage groups were identified: alveolar macrophages (fatty acid binding protein 4 (*FABP4*)+), interstitial macrophages (legumain (*LGMN*)+), monocytes/recently-differentiated macrophages (ficolin-1 (*FCN1*)+) and proliferating macrophages (*MKI67*+), based on canonical marker expression (Figure S3A-B). Comparative analysis revealed significantly higher proportion of proliferating macrophages in RAS lungs compared to controls, with a trend towards increased IMΦs relative to BOS (Figure S3C-D). The proportion of monocyte-derived macrophages was similar across conditions.

For higher-resolution characterization, the dataset was reclustered using unbiased Louvain clustering at an optimized resolution of 1.0, as determined by Clustree analysis (24) (Figure S3E), yielding 12 distinct macrophage subsets, which were annotated based on differential gene expression and prior literature (Figure 1C; Table 2). These included interferon-alpha inducible protein 27 (*IFI27*) (25) and fibronectin 1 (*FN1)+* alveolar macrophages (26, 27)); folate receptor beta (*FOLR2)*+ (28, 29) and coagulation factor XIII A chain (*F13A1)*+ interstitial macrophages (28, 29), interferon-stimulated gene (ISG) macrophages (22, 30), *SPP1* macrophages (4, 31)), monocytes (*CD14*+, *S100A8/A9*+) (30), *CXCL8*+ monocytes (*CD14*+ *S100A8/A9*+ *CXCL8*+) (32), galectin-2 (*LGALS2*)+ monocytes (33, 34), LILR macrophages (35), CD301 (*CLEC10A)*+ macrophages (36)) and proliferating macrophages (*MKI67*+ *TOPA2*+) (25, 37).

**Table 2.** Functional Annotation of Macrophage Subsets.

| Cluster name | Specialty/Function | Reference |
| --- | --- | --- |
| IFI27+ AMs | Viral defense. | Li et al., 2022 (25) |
| FN1+ AMs | M2-like AM | Jablonski et al., 2015 (26);<br>Huang et al., 2019 (27); |
| FOLR2+ IMs | CD169+ IMs, conserved TLF+ macrophages | Hume et al., 2025 (28); Dick et al., 2022 (29) |
| Monocytes | Classical monocytes | Mulder et al., 2021 (30) |
| CLEC10A+ | M2-like monocytes | Browning et al., 2025 (36) |
| ISGs | Interferon-stimulated macrophages | Moshkelgosha et al., 2021 (22); Mulder et al., 2021 (30) |
| Galectin-2 | M1-like monocyte | Shi et al., 2025 (33);<br>Yıldırım et al., 2015 (34) |
| LILR | Leukocyte immunoglobulin-like receptor enriched macrophages | Varin et al., 2025 (35) |
| CXCL8+ Monocytes | Pro-inflammatory monocytes | MacDonald et al., 2021 (32) |
| Proliferating | Proliferating macrophages | Li et al., 2022 (25); Wu et al., 2023 (37) |
| F13A1+ IMs | Antigen presentation | Hume et al., 2025 (28); Dick et al., 2022 (29) |
| SPP1 | Pro-fibrotic macrophages | Morse et al., 2019 (4) ;<br>Reyfman et al., 2019 (31) |

### Distinct macrophage subsets are differentially enriched in BOS and RAS

Comparison of the relative abundance of the 12 macrophage subsets revealed several populations enriched in CLAD lungs, including ISG, LILR, proliferating and SPP1 macrophages (Figure 1D, Figure S3F). Enrichment of ISG and proliferating macrophages in CLAD lungs were consistent with prior reports (22, 38). When stratified by CLAD phenotype, LILR macrophages were significantly enriched in BOS, whereas SPP1 macrophages were significantly enriched in RAS (Figure 1E). To define transcriptional differences associated with disease, differential gene expression analysis using model-based analysis of single-cell transcriptomics (MAST) (39) identified genes enriched in CLAD macrophages compared to control lungs (Figure 1F). Further comparisons between BOS and RAS macrophages demonstrated that pro-fibrotic genes, including *SPP1*, *MMP9* and *CHI3L1*, were preferentially associated with RAS phenotype (Figure 1G).

### LILR and SPP1 macrophages exhibit pro-inflammatory and pro-fibrotic transcriptomic programs, repectively

Differential gene expression analysis identified distinct transcriptional programs in LILR and SPP1 macrophage subsets (Figure S4). LILR macrophages preferentially expressed genes associated with immune activation, including *CD300E*, *APOBEC3A*, and *LILRA* family members, whereas SPP1 macrophages were enriched for pro-fibrotic genes such as *MMP9*, *CHI3L1*, *COL6A2*, *MMP7*.

To validate these transcriptional signatures, inflammatory and fibrotic gene module scores were calculated using curated gene sets from published pulmonary macrophage-focused scRNAseq studies of type I inflammation and idiopathic pulmonary fibrosis (IPF) (Supplemental Table 3) (40–43). LILR macrophages exhibited significantly elevated inflammatory gene module scores relative to non-LILR macrophage populations (Figure 2A; Figure S5A), as it also demonstrated by increased expression of selected pro-inflammatory genes (*LILRA5*, *IFITM2* and *ISG20*; Figure 2B). Consistent with enrichment of phagocytosis-associated transcripts, LILR macrophages also demonstrated elevated phagocytic and engulfment gene module scores (44, 45) compared to other subsets (Figure S5B-C). Gene Ontology enrichment analysis further identified immune-associated pathways, including lymphocyte activation and cell adhesion, among the top enriched biological processes (Figure 2C).

**Figure 2:**
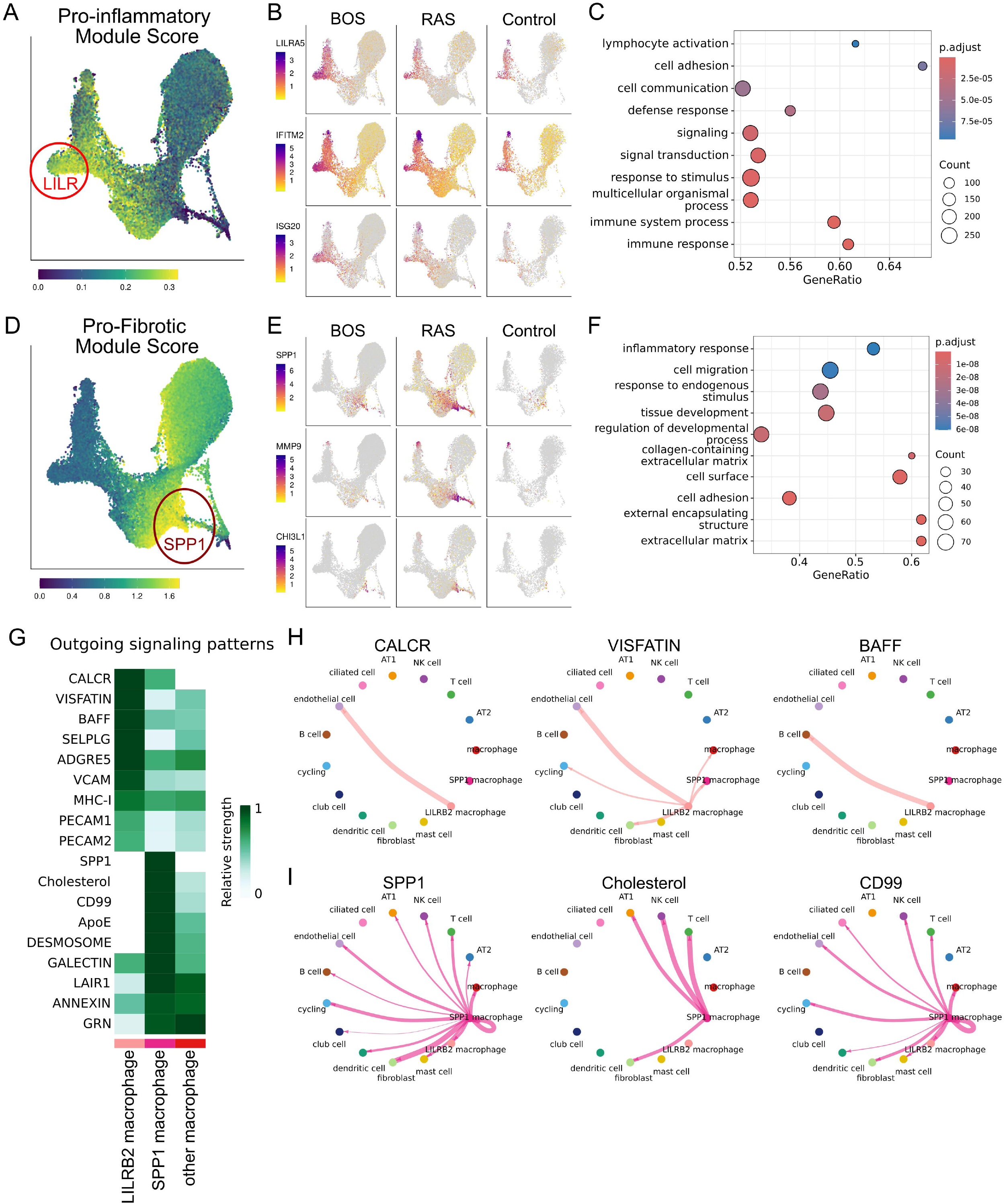
Pro-inflammatory and pro-fibrotic transcriptional programs and outgoing signaling interactions of LILR and SPP1 macrophages. Single-cell RNA sequencing data from 14 CLAD lungs (BOS = 7, RAS = 7) and 6 control lungs is shown. A) UMAP visualization of enrichment scores for pro-inflammatory gene programs. B) Feature plots showing relative expression of representative pro-inflammatory genes: LILRA5, IFITM2 and ISG20. C) Gene set enrichment analysis dot plot showing the top 10 GO biological processes enriched in LILR macrophages. D) UMAP visualization of enrichment scores for pro-fibrotic gene programs. E) Feature plots showing relative expression of representative pro-fibrotic genes: SPP1, MMP9, and CHI3L1. F) Gene set enrichment analysis dot plot showing the top 10 GO biological processes enriched in SPP1 macrophages. G) Heatmap of top outgoing signaling pathways from LILR and SPP1 macrophages inferred by CellChat, showing relative interaction strength across macrophage subsets. H) Circle plot illustrating LILR macrophage interactions through dominant outgoing pathways (CALCR, VISFATIN, BAFF). I) Circle plot illustrating SPP1 macrophage interactions through dominant outgoing pathways (SPP1, Cholesterol, CD99).

The SPP1 macrophage cluster exhibited significantly higher fibrotic gene module score (Figure 2D; Figure S5D) and strong expression of canonical pro-fibrotic genes, with preferential enrichment in RAS lungs (Figure 2E). Gene Ontology enrichment analysis revealed elevated enrichment in inflammatory response and multiple extracellular matrix remodeling related pathways within this cluster (Figure 2F).

Collectively, these findings demonstrate that LILR macrophages exhibit a dominant pro-inflammatory transcriptional program, whereas SPP1 macrophages are characterized by a pro-fibrotic, tissue-remodeling gene expression profile, supported by differential gene expression, gene module scoring and pathway-level enrichment analysis.

### LILR and SPP1 macrophage states exhibit divergent transcription factor programs

To further validate inflammatory and fibrotic macrophage identities at the regulatory level, SCENIC analysis was performed to infer transcription factor regulon activity across macrophage subsets (46). LILR macrophages demonstrated enriched activity of regulons associated with inflammatory transcriptional remodeling, including ZBTB7A (47), IKZF1 (48), and BCLAF1 (49), along with regulatory programs linked to immune modulation (KLF2 (50) and TCF7L2 (51)) (Figure S5E). In contrast, SPP1 macrophages exhibited increased activity with ARID5B, MITF, and GRHPR regulons, which collectively are involved with tissue remodeling and metabolic adaptation (Figure S5E) (52–54). Together, these regulon profiles support a pro-inflammatory regulatory program in LILR macrophages and a pro-fibrotic, tissue-remodeling program in SPP1 macrophages, consistent with differential gene expression, module scoring and pathway-level analysis.

### Inferred cell-cell communication reveals distinct interaction networks for LILR and SPP1 macrophages

To identify potential downstream targets of LILR and SPP1 macrophages, ligand-receptor interactions were inferred using CellChat analysis (55). The top outgoing signaling pathways from the LILR and SPP1 macrophages were ranked and compared with other macrophage subsets (Figure 2G). LILR macrophages exhibited strongest signaling through pathways associated with inflammation, such as the calcitonin receptor pathway (CALCR) (56) and nicotinamide phosphoribosyltransferase (NAMPT)/visfatin (VISFATIN) pathway (57, 58), or interactions with lymphoid cells such as B-cell activating factor pathway (BAFF), whereas SPP1 macrophages showed dominant activity in SPP1, cholesterol and CD99 pathways, which are all involved in fibrosis and tissue remodeling (59–62).

Analysis of inferred target cell populations revealed that CALCR and VISFATIN signaling from LILR were primarily directed towards endothelial cells, while BAFF signaling targeted B cells (Figure 2H). In contrast, SPP1 macrophage signaling via the SPP1 pathway was broadly distributed but predominantly weighted towards fibroblasts. Cholesterol signaling targeted alveolar type I cells, NK cells, T cells and fibroblasts, whereas CD99 signaling was directed towards most cell types (Figure 2I). Transcript-level expression of key ligand-receptor pairs underlying these pathways was confirmed across relevant clusters (Figure S6A-B). Together, these analyses suggest that LILR and SPP1 macrophages may engage in distinct ligand-receptor communication programs with immune, endothelial and stromal cell populations, consistent with their transcriptionally defined inflammatory and fibrotic roles.

### Independent CLAD datasets confirm enrichment of LILR macrophages

To assess the reproducibility of the LILR and SPP1 macrophage populations, we first examined previously published CLAD scRNAseq datasets from Khatri et al. (63) and Yan et al. (38) (Duke/Northwestern). The Duke/Northwestern dataset contained broadly similar immune and parenchymal cell populations to our Toronto cohort, although neutrophils were not observed in the Toronto cohort (Figure S7A). Reclustering of macrophages identified a distinct LILR macrophage population in the Duke/Northwestern dataset, whereas SPP1 macrophages were not detected (Figure S7B). Integration of macrophages from the Duke/Northwestern and Toronto cohorts demonstrated that LILR macrophages were consistently represented across datasets, whereas SPP1 macrophages were only observed in the Toronto cohort (Figure S7C-D), potentially reflecting cohort-specific differences, including the limited representation of RAS in the Duke/Northwestern dataset.

### Comparison with IPF lungs reveals conserved fibrotic macrophage programs

Given the absence of SPP1 macrophages in public CLAD datasets, we next examined whether LILR and SPP1 macrophage populations were conserved in another fibrotic lung disease. Two publicly available IPF scRNAseq datasets (GSE122960, GSE128033) were analyzed independently of our CLAD cohort and annotated (Figure S7E). IPF macrophages were isolated and reclustered, revealing distinct LILR and SPP1 macrophage populations based on expression of canonical marker genes (Figure S7F). Macrophages from the IPF datasets were then integrated with CLAD macrophages, revealing transcriptional concordance between LILR and SPP1 populations across disease states (Figure S7G-H). Notably, LILR-associated genes were markedly higher in CLAD macrophages compared to IPF, suggesting amplification of inflammatory macrophage programs in CLAD, potentially reflecting the alloimmune milieu inherent to CLAD but absent in IPF.

### Cross-tissue analysis positions LILR and SPP1 macrophages within conserved myeloid programs

To contextualize pulmonary macrophage subsets within broader myeloid biology, we compared our data with a published cross-tissue single-cell atlas of human monocytes and macrophages spanning multiple tissues and disease states (30). Differentially expressed gene signatures from atlas-defined populations were compared with macrophage subsets identified in our dataset to assess transcriptional concordance (Figure S8A).

Six atlas-defined populations showed transcriptional similarity to macrophage subsets in our study. Trem2 macrophages showed the strongest similarity to SPP1 macrophages, whereas C1Qhi macrophages mapped to a subset of the alveolar macrophage population. Their proliferating macrophages showed similarities to our proliferating macrophage population, IL-4I1 macrophages to ISG macrophages, CD16^+^ monocytes to LILR macrophages, and CD16^-^ monocytes to our annotated monocytes. Notably, although SPP1 macrophages showed similarity to TREM2 tumor-associated macrophages (TAMs), the SPP1 population identified here exhibited robust expression of pro-fibrotic genes not typically associated with TAMs. This observation is consistent with the presence of heterogeneous SPP1-expressing macrophage states, potentially encompassing tumor-associated and fibrosis-associated programs (64).

LILR macrophage additionally demonstrated transcriptional similarity to CD16^+^ monocytes, consistent with a potential non-classical monocytic origin. Consistent with the limitations of classical M1/M2 framework in describing in vivo macrophage states, neither LILR or SPP1 macrophages aligned exclusively with M1 or M2 polarization signatures, and both populations exhibited overlapping M1 and M2 gene module scores (Figure S8B). Interstitial macrophages were less clearly represented in this atlas, although FOLR2+ IMs and CLEC10A+ macrophages shared features with tissue-resident macrophages described in independent cross-tissue analyses (29). Together, cross-disease, cross-cohort and cross-tissue analyses support the robustness and biological relevance of distinct inflammatory LILR and fibrotic SPP1 macrophage programs identified in CLAD.

### SDC and LILRB2 enable isolation of pro-fibrotic and pro-inflammatory macrophages

To enable functional characterization of transcriptomically defined LILR and SPP1 macrophage populations, we used Cerebro, an R package that identifies differentially expressed genes encoding cell surface proteins (65) to identify candidate markers suitable for their isolation. *LILRA5* and *LILRB2* were identified as highly enriched in the LILR macrophages, with their expression predominantly localized to this population based on feature plots (Figure 3A-B). Allograft inflammatory factor 1 (*AIF1*), a non-surface macrophage marker, was also preferentially expressed in LILR macrophages. Although not suitable for surface-based isolation, AIF1 was included as a complementary transcriptional marker for downstream validation by RT-qPCR. For SPP1 macrophages, syndecan-2 (*SDC2*), *CD36*, and integrin-β5 (*ITGB5*) were identified as candidate surface markers; however, only *SDC2* demonstrated preferential expression specific to the SPP1 cluster whereas *CD36* and *ITGB5* were broadly expressed (Figure 3C-D). SDC2 was therefore selected as the optimal surface marker for isolating SPP1 macrophages.

**Figure 3:**
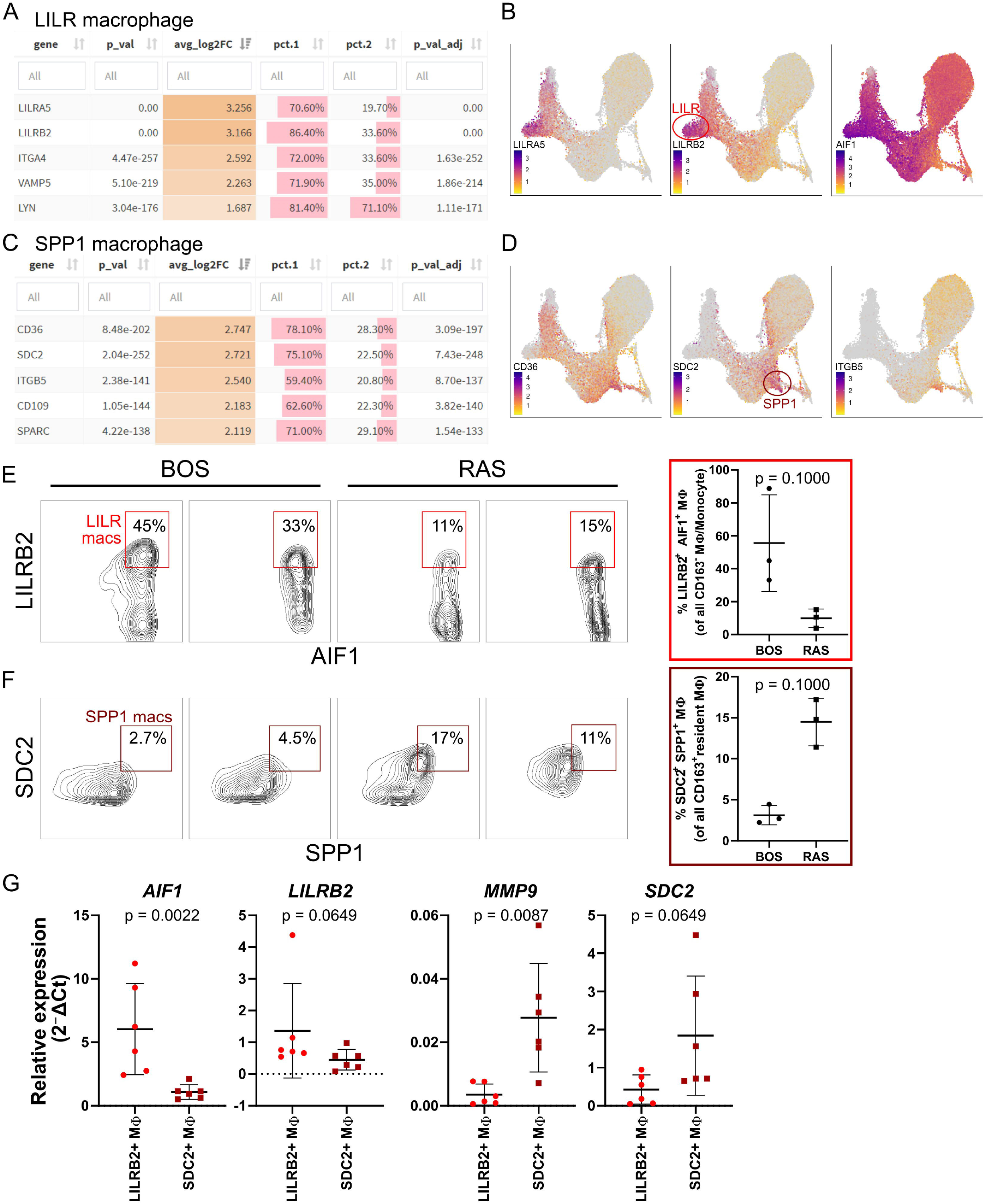
Surface marker–based isolation and validation of LILR and SPP1 macrophage populations. A) Top 5 differentially expressed genes encoding surface marker proteins for LILR macrophages in single-cell RNA sequencing data is shown. B) Feature plots showing expression of top two surface-protein-encoding genes (LILRA5 and LILRB2), along with AIF, across lung macrophages. C) Top 5 differentially expressed genes encoding surface marker proteins for SPP1 macrophages in single-cell RNA sequencing is shown. D) Feature plots showing expression of top three surface-protein-encoding genes (CD36, SDC2, ITGB5). E) Representative flow cytometry plots from BOS and RAS lung samples showing AIF1 x LILRB2 expression from gated CD163^-^ macrophage/monocytes (left) and quantification of double positive macrophages as a percentage of CD163^-^ macrophage/monocytes, stratified by BOS and RAS (right). F) Representative flow cytometry plots from BOS and RAS lung samples showing SPP1 x SDC2 expression from gated CD163^+^ resident macrophages (left) and quantification of double positive macrophages as a percentage of CD163^+^ resident macrophages, stratified by BOS and RAS (right). G) Relative transcript expression of selected markers (*AIF1*, *LILRB2*, *MMP9*, *SDC2*) in sorted LILRB2+ and SDC2+ macrophages, measured by RT-qPCR and expressed as 2^-ΔCt^. For panels E-G, statistical comparisons were performed using a Mann-Whitney U test, with p-values indicated in the figure.

Protein-level specificity was validated by flow cytometry of CLAD lung single-cell suspensions, following gating on CD14^+^CD163^-^ monocyte/macrophages and CD14^+^CD163^+^ resident macrophages (Figure S9A). Within the monocyte/macrophage compartment, LILRB2 and AIF1 co-expression was significantly increased in BOS samples (Figure 3E), whereas SDC2 and SPP1 co-expression was enriched in CD14^+^CD163^+^ resident macrophages from RAS lungs (Figure 3F), consistent with scRNAseq distributions.

To further confirm concordance between sorted populations and transcriptionally defined clusters, fluorescence-activated cell sorting (FACS) was performed on cryopreserved CLAD lung samples derived from scRNAseq cases, followed by reverse transcription-quantitative polymerase chain reaction (RT-qPCR) validation (Figure S9A-B). AIF1 mRNA expression was significantly higher in LILRB2+ macrophages, with LILRB2 transcript showing a similar trend, whereas MMP9 mRNA expression was significantly enriched in SDC2+ macrophages, along with a trend towards increased SDC2 transcript expression (Figure 3G). Collectively, concordant findings from scRNAseq, flow cytometry and RT-qPCR demonstrate that LILRB2 and SDC2 reliably identify and enable isolation of pro-inflammatory LILR and pro-inflammatory SPP1 macrophage populations, respectively.

### LILR macrophages exhibit enhanced phagocytic capacity and pro-inflammatory effector functions

To functionally validate the pro-inflammatory phenotype inferred from transcriptomic analyses, LILR, SPP1 and SDC2^-^ resident macrophages (SDC2^-^LILRB2^-^ macrophages) were isolated by spectral-enhanced FACS from cryopreserved CLAD lung samples (n = 12; 6 BOS, 6 RAS; Figure S9C). This yielded 12 biological replicates of LILR macrophages, 9 biological replicates of SPP1 macrophages and 12 biological replicates of SDC2^-^LILRB2^-^ macrophages for downstream *in vitro* assays. Three BOS lungs did not yield sufficient SPP1 macrophages for downstream analyses. Patient characteristics of these patients are reported in Table 3.

**Table 3.**
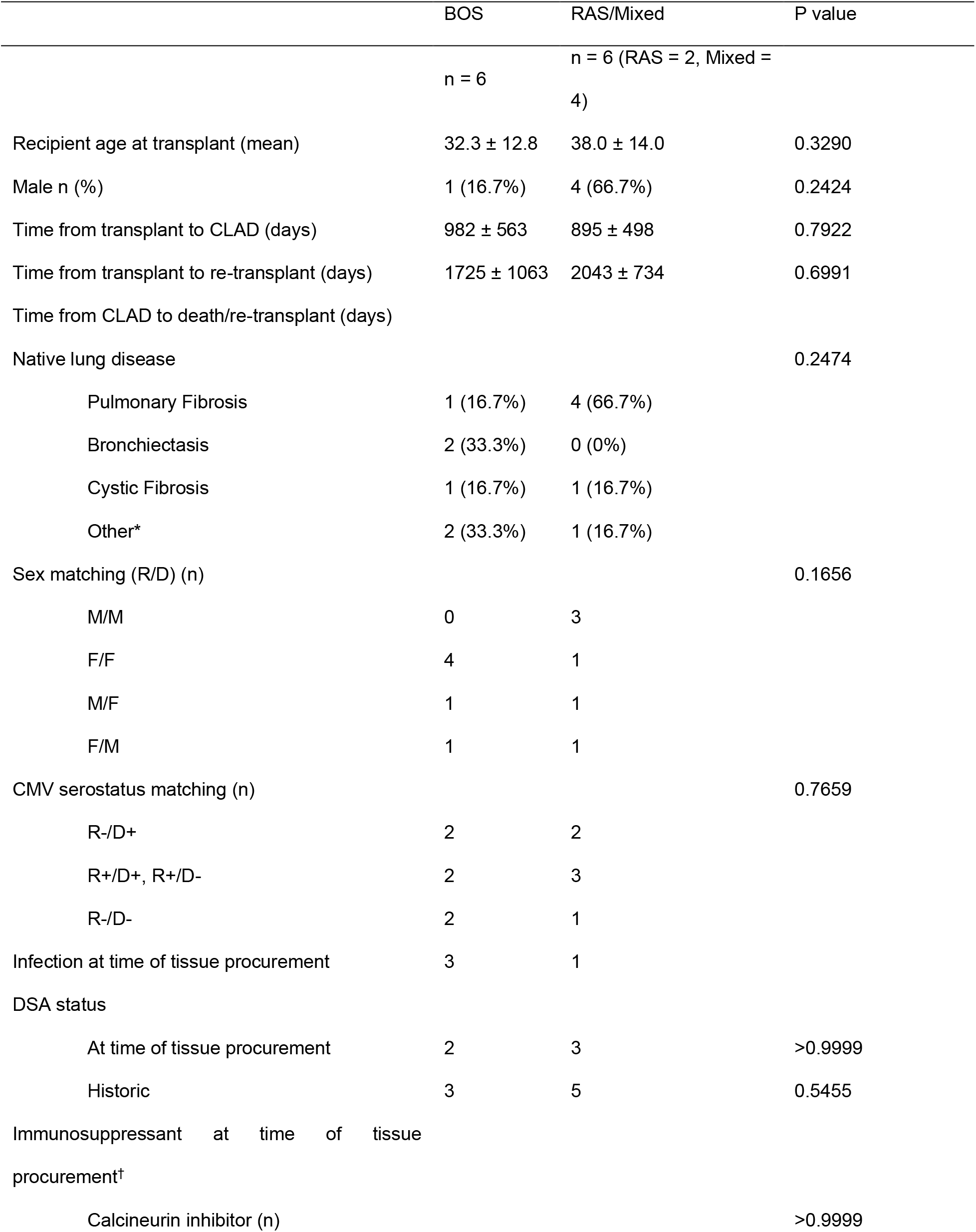
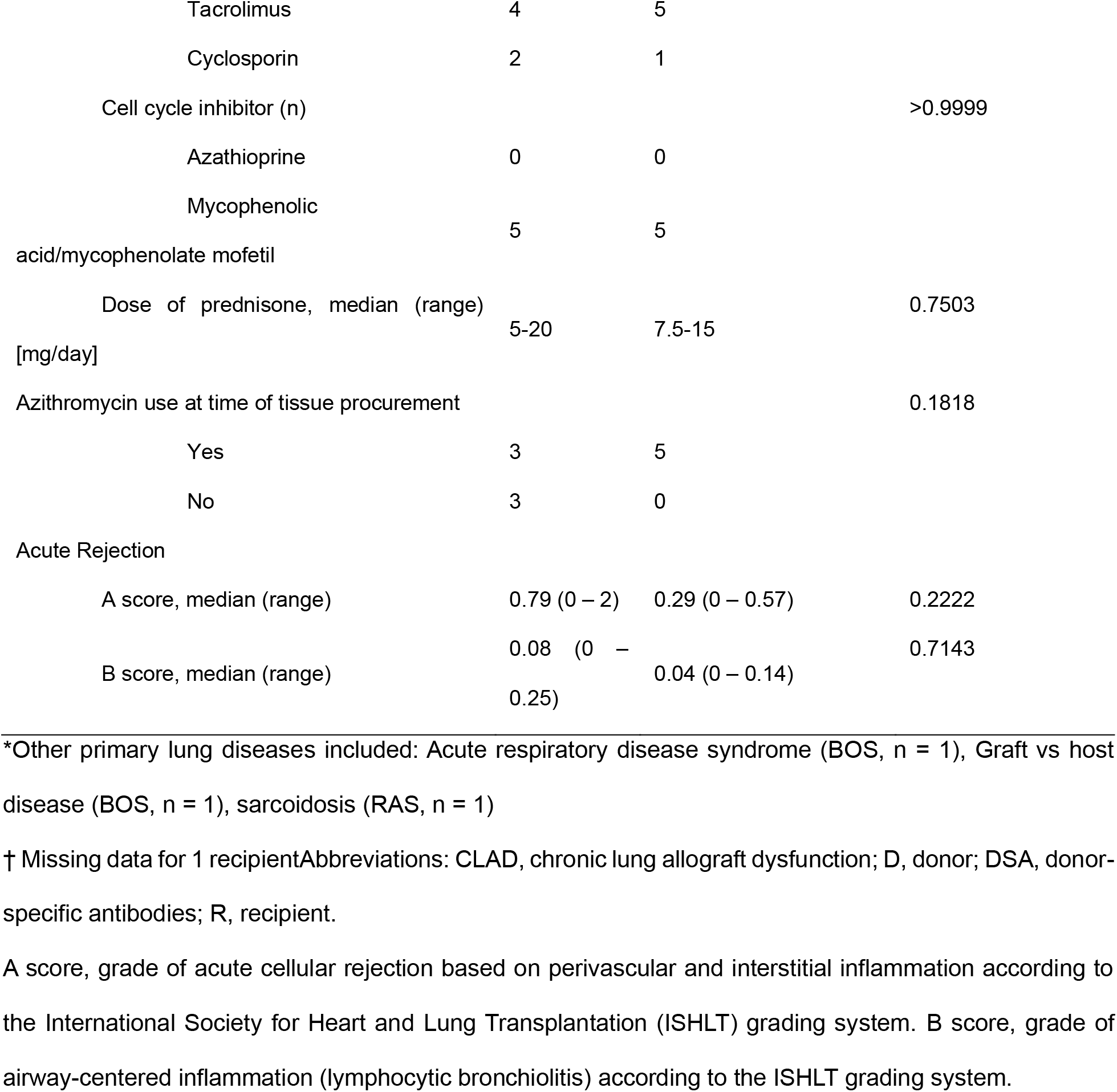
Sorted CLAD Cohort Patient Characteristics.

Consistent with enrichment of phagocytosis-associated transcripts in LILR macrophages, phagocytic activity, assessed by pHrodo Red *E. coli* phagocytosis assay, demonstrated significantly greater *E. coli* uptake in LILR macrophages compared with SPP1 and SDC2^-^ LILRB2^-^ (Figure 4A-B). Given transcriptomic enrichment of lymphocyte-interaction pathways, the ability of supernatants from LILR macrophages – as compared to those from SPP1 and SDC2^-^LILRB2^-^ macrophages – to promote T cell chemotaxis was assessed using transwell migration assays with macrophage-conditioned supernatant (Figure 4C). Activated T cells migrated preferentially toward supernatant from LILR macrophages compared with SPP or SDC2^-^LILRB2^-^ macrophages supernatants (Figure 4D).

**Figure 4:**
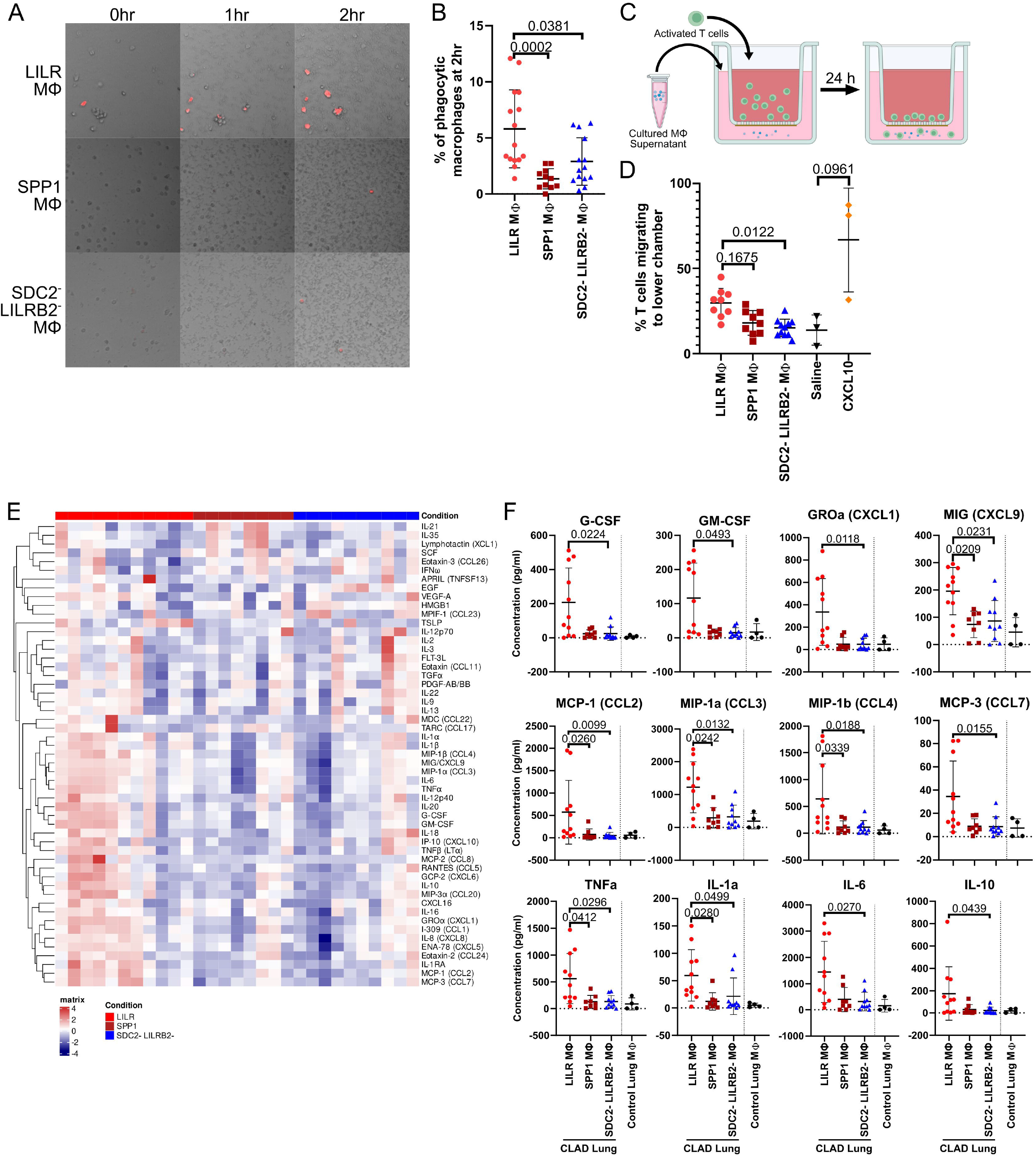
LILR macrophages demonstrate pro-inflammatory functions *in vitro*. A) Representative micrographs showing phagocytosis of pHrodo Red™ *E. coli* bioparticles by cultured macrophages (MΦ) obtained from CLAD lungs, over time. Images were acquired with a 10x objective lens. B) Quantification of phagocytic macrophages across LILR, SPP1, and SDC2^-^LILRB2^-^ MΦ, with individual biological replicates shown. C) Schematic diagram of the T cell chemotaxis assay, showing migration of activated T cells from the upper chamber towards conditioned media placed in the lower chamber. D) Quantification of T cell migration towards conditioned media from LILR, SPP1, or SDC2^-^ LILRB2^-^ MΦ, as well as saline and CXCL10 controls, with individual biological replicates shown. E) Heatmap showing relative concentrations of soluble analytes from multiplex assay, across LILR, SPP1 and SDC2^-^LILRB2^-^ MΦ populations. Macrophage subsets were grouped by population, whereas analytes were unsupervised and clustered based on their concentration profiles. F) Quantification of concentrations of analytes significantly higher in LILR MΦ supernatants, compared to SPP1 and SDC2^-^LILRB2^-^ MΦ populations, with individual biological replicates shown. Supernatants from non-CLAD (excess donor lung) samples were included for reference as a baseline but were not included in statistical comparisons. For panels B, D and F, group comparisons were performed using a Kruskal-Wallis test, with post-hoc multiple comparisons performed; corresponding P values are shown.

Based on inferred ligand-receptor interactions with endothelial cells, endothelial cell activation assays were performed using cultured endothelial cells exposed to macrophage-conditioned supernatants (Figure S10A). No significant changes in endothelial activation markers were observed relative to saline controls (Figure S10B), suggesting that LILR macrophage-endothelial interactions are unlikely to be mediated by soluble factors.

To define soluble mediators contributing to T cell chemotaxis and identify additional pro-inflammatory mediators, conditioned supernatants were analyzed using multiplex bead-based cytokine and chemokine profiling. Compared with SPP1 and SDC2^-^LILRB2^-^ macrophages, LILR macrophages exhibited a selectively pro-inflammatory secretory profile (Figure 4E), with significantly increased pro-inflammatory cytokines, and chemokines (Figure 4F). Several additional analytes were upregulated from LILR macrophages but did not reach significance (Figure S11A), whereas others were unchanged across macrophage subsets (Figure S11B).

Collectively, these results demonstrate that LILR macrophages exhibit pro-inflammatory functional activity *in vitro*, including enhanced phagocytosis, T cell chemotaxis and production of inflammatory cytokines and chemokines. These functions are consistent with their transcriptional profile and supporting a role for LILR macrophages in driving inflammatory responses in CLAD.

### SPP1 macrophages induce a pro-fibrotic fibroblast phenotype

Given the inferred interactions between SPP1 macrophages and fibroblasts, we assessed the functional impact of SPP1 macrophage-derived soluble factors on fibroblast behavior. MRC-5 fibroblasts were cultured with conditioned supernatants from SPP1, LILR or SDC2^-^LILRB2^-^ macrophages for three days. Controls were normally migrating fibroblasts cultured with medium plus saline and fibroblasts treated with TGF-β to reduce migration (66). All fibroblast monolayers were then scratched to create a gap and their migratory capacity closing the gap was monitored over 24 hours (Figure 5A). Fibroblasts treated with SPP1 macrophage supernatants exhibited significantly reduced migration at 24 hours compared with fibroblasts treated with SDC2^-^LILRB2^-^ macrophage supernatants (Figure 5B). Although SPP1- and LILR-treated fibroblasts showed reduced migration relative to saline controls and were comparable to TGF-β-treated controls, SPP1 macrophage supernatant induced the most pronounced impairment of migration. Next, MRC-5 fibroblasts were embedded in attached collagen gels to measure their contractile activity as another function acquired during their activation (67) (Figure 5C). Fibroblasts treated with SPP1 macrophage supernatants exhibited the greatest collagen gel contraction, contracting by 35% within90 minutes after detaching the collagen, comparable to that of TGF-β-treated positive controls (Figure 5D). In contrast, LILR macrophage supernatant-treated fibroblasts showed minimal contraction (20%), similar to saline treated controls.

**Figure 5:**
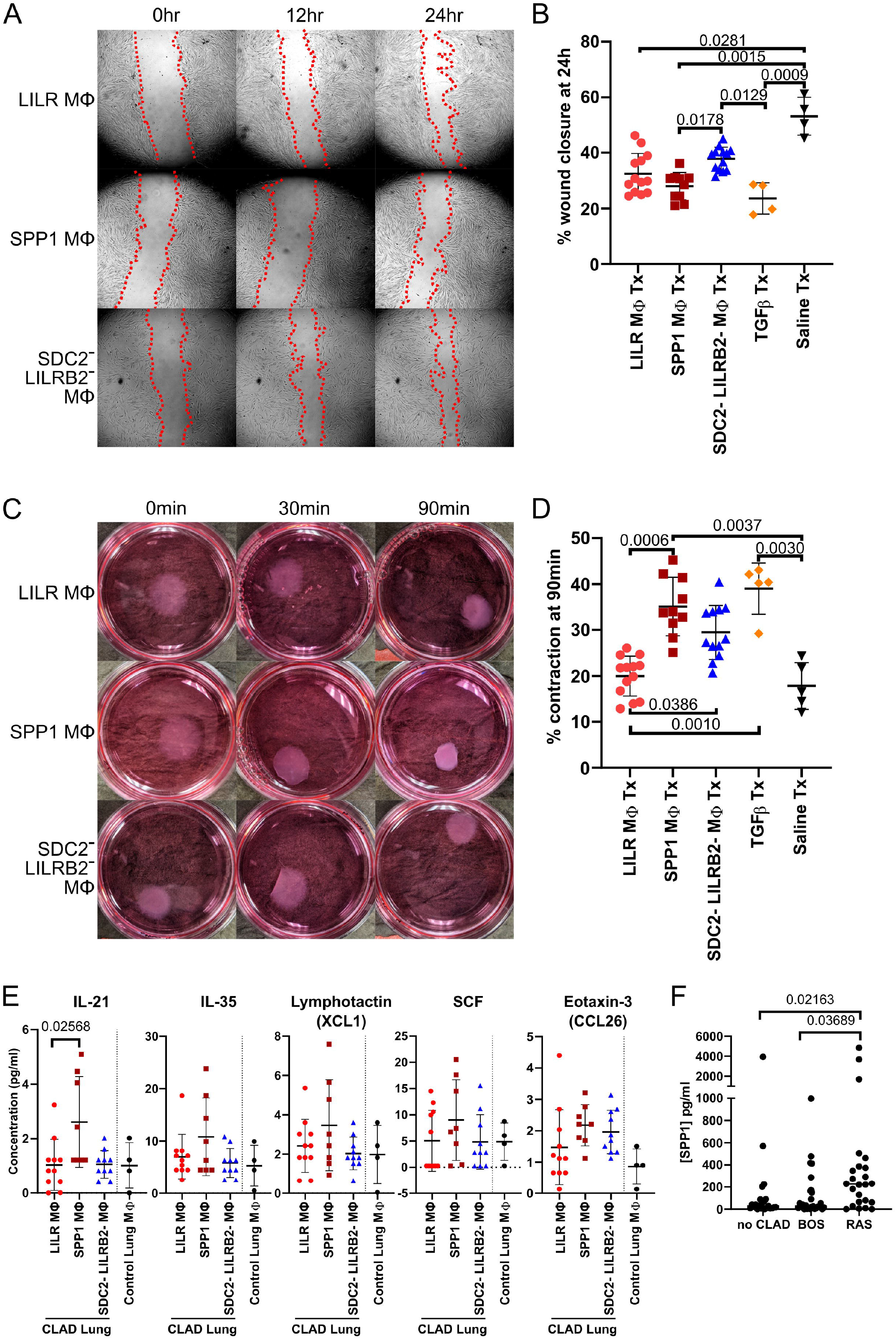
SPP1 macrophages promote fibroblast migratory and contractile responses *in vitro*. A) Representative micrographs from a scratch (wound-healing) assay showing fibroblast migration in response to conditioned media from LILR, SPP1, and SDC2^-^LILRB2^-^ macrophage (MΦ) populations obtained from CLAD lungs. Images are shown at 0, 1, and 2 hours following scratch generation. B) Quantification of wound closure at 2 hours following exposure to macrophage-conditioned media, with saline and TGF-β-treated conditions included as negative and positive controls. Individual biological replicates are shown. C) Representative images from a collagen gel contraction assay showing fibroblast-mediated gel contraction over time (0, 30, and 90min) following exposure to macrophage-conditioned media. D) Quantification of collagen gel contraction at 90 minutes across macrophage-conditioned media and control conditions, with individual biological replicates shown. E) Quantification of concentrations of analytes significantly higher in SPP1 MΦ supernatants, compared with LILR and SDC2^-^LILRB2^-^ MΦ populations, with individual biological replicates shown. Supernatants from non-CLAD (excess donor) lung samples were included for references as a baseline but were not included in statistical comparisons. F) Quantification of soluble SPP1 in BAL from lung transplant recipients without CLAD, with BOS, or with RAS/mixed CLAD phenotypes. For panels B, D-F, group comparisons were performed using a Kruskal-Wallis test with post-hoc multiple comparisons testing; corresponding adjusted P values are shown.

To identify candidate mediators underlying these pro-fibrotic effects, we re-examined the multiplex cytokine profiling data from macrophage-conditioned supernatants, focusing on analytes enriched in SPP1 macrophages (Figure 4E). Several mediators showed higher mean concentrations in SPP1 macrophage supernatant, with interleukin (IL)-21 reaching statistical significance (Figure 5E); IL-21 is known to induce STAT3 phosphorylation in fibroblasts (68). As SPP1 was not included in the multiplex panel, a separate SPP1-specific ELISA was performed on bronchoalveolar lavage samples from a cohort of lung transplant recipients without CLAD, with BOS-type CLAD, or with RAS/mixed-type CLAD from a previous study (69). SPP1 protein levels were significantly elevated in BAL from patients with RAS/mixed compared to no-CLAD and BOS groups (Figure 5F). Collectively, these data demonstrate that SPP1 macrophages secrete soluble mediators that promote fibroblast migration arrest and enhanced contractility, consistent with a myofibroblast-like, pro-fibrotic phenotype.

## DISCUSSION

In this study, we identified two macrophage subsets enriched in CLAD: pro-inflammatory LILR and pro-fibrotic SPP1 macrophages. These populations exhibited distinct transcriptional programs associated with inflammation and fibrotic remodeling, respectively, suggesting specialization in divergent pathogenic processes within the allograft. Using sorting strategies based on differentially expressed surface markers, we isolated LILR and SPP1 macrophages and directly confirmed their pro-inflammatory and pro-fibrotic functions using complementary *in vitro* assays.

LILR macrophages, characterized by a pro-inflammatory transcriptional profile, are likely recently monocyte-derived, based on their shared transcriptional signatures with monocyte populations in our dataset and in independent public datasets (30). Further studies will be needed to validate their localization; however, their ability to recruit circulating immune cells and amplify inflammatory responses within the allograft positions LILR macrophages as a key pathogenic population in CLAD. Additional studies will also be required to clarify the inferred interactions between LILR macrophages and endothelial cells, which may require direct cell-to-cell contact. While LILR-expressing macrophages have not been extensively described in prior studies of lung disease, they have been implicated in promoting alloimmune T cell responses in kidney transplantation, annotated as LILRB2+ CXCL10+ macrophages (35).

Inflammatory macrophage subsets have been described across CLAD and other lung diseases using single-cell approaches. ISG macrophages have been previously identified in CLAD (22, 38), and are characterized by a pro-inflammatory, interferon-γ (IFNγ)-responsive gene signature with expression of chemokines CXCL10 and CXCL11. These populations appear distinct from LILR macrophages, as evidenced by their separate clustering and divergent transcriptomic profiles. However, both ISG and LILR macrophages appear to be linked through IFNγ signaling, as IFITM2 and ISG20, and other IFNγ responsive genes are differentially expressed in LILR macrophages. This suggests that, while these macrophage subsets may be driven by common upstream signals, they adopt distinct and complementary effector roles that contribute to inflammation and alloimmune responses in CLAD.

In contrast to this inflammatory macrophage program, SPP1 macrophages were selectively enriched in RAS in our dataset and exhibited a pro-fibrotic transcriptional and functional profile. SPP1 macrophage-conditioned supernatants reduced fibroblast migration and increased contractility, consistent with myofibroblast activation (70). Cytokine profiling revealed increased levels of IL-21, IL-35, CCL26 and XCL1, all of which have been associated with pulmonary fibrogenesis (71–74). Consistent with these *in vitro* findings, soluble SPP1 protein was significantly elevated in BAL samples from CLAD patients with the RAS phenotype, which our scRNAseq analysis indicated was predominantly produced by SPP1 macrophages. Through signaling via fibroblast receptors such as CD44 and integrin β3, SPP1 can promotes proteolytic activation of latent TGF-β and subsequent engagement of the TGF-β/SMAD signaling pathway (61, 75). Activation of this pathway alters fibroblast transcriptional programs, driving myofibroblast differentiation and pathologic extracellular matrix remodeling, situating SPP1 macrophages as plausible cellular drivers of fibrotic injury in RAS-type CLAD.

SPP1 macrophages were not detected in appreciable numbers in BOS samples in our dataset nor in other CLAD scRNAseq studies (22, 38, 63), suggesting that the small-airway-predominant fibrosis characteristic of BOS may arise through alternative fibrotic pathways or cell types. In contrast, fibrotic interstitial lung disease such as IPF show consistent enrichment of SPP1 macrophages within fibrotic niches (4, 31, 76), a finding we confirmed through integration of CLAD and public IPF datasets (4, 31). SPP1 macrophages have also been identified across multiple organs as key contributors to fibrotic remodeling, underscoring the importance of this population in fibrotic disease biology (75, 77–79). Together, these observations underscore the central role of SPP1 macrophages in fibrotic remodeling and highlight the importance of their isolation for mechanistic and therapeutic studies in CLAD and other fibrotic diseases.

These findings have important clinical and translational implications for CLAD. Our data suggest macrophage-based mechanisms that may enable earlier phenotypic stratification of CLAD and provide targets directed at the inflammatory and fibrotic processes that drive CLAD progression. Given their role in promoting alloimmune responses, LILR macrophages are a promising therapeutic target; strategies aimed at modulating LILR-mediated signaling may offer a means to attenuate macrophage-driven inflammation in CLAD (35, 80). Through their direct effects on fibroblast activation and matrix remodeling, SPP1 macrophages represent a therapeutic focus in RAS-type CLAD through the SPP1-dependent fibrotic signaling. Prior studies demonstrating attenuation of fibrosis following inhibition of SPP1 signaling further support the therapeutic relevance of this pathway (62). In addition, the selective enrichment of SPP1 in RAS lung tissue and detection of soluble SPP1 in BAL samples raise the possibility that macrophage-based biomarkers could enable earlier identification of high-risk CLAD phenotypes and permit timely antifibrotic treatment.

This study has several limitations. First, lung tissue samples were obtained exclusively at end-stage CLAD, which does not capture earlier disease processes. As such, whether LILR and SPP1 macrophage populations arise earlier in CLAD progression remains to be determined. Second, the sample size was modest, reflecting the technical and logistical challenges of performing scRNAseq on explanted CLAD tissue. Despite this limitation, key cell populations, including LILR and SPP1 macrophages were consistently identified across samples, supporting the robustness of our findings. Third, *in vitro* functional assays may not fully recapitulate the complexity of *in vivo* behavior. Nonetheless, these experiments provided mechanistic insight concordant with scRNAseq-derived signatures.

In summary, this study identifies LILR and SPP1 macrophages as distinct pathogenic subsets in CLAD and defines their inflammatory and fibrotic functional properties through integrated single-cell transcriptomic and targeted *in vitro* analyses. Beyond mechanistic insight into the potential roles of specific macrophage populations in allograft injury, our findings establish sorting strategies that enable isolation and functional interrogation of these populations in CLAD and with potential applicability to other transplant and fibrotic disease contexts. Together, these results provide a framework for future studies to examine macrophage programming at earlier stages of CLAD and to explore macrophage-targeted therapeutic approaches aimed at modulating inflammatory and fibrotic progression.

## METHODS

### Sex as a biological variable

Both male and female participants were included in this study. The study was not designed or powered to evaluate sex-specific differences, and sex was therefore not incorporated as a biological variable in the primary analyses. Macrophage populations and functional responses were analyzed irrespective of sex, although sex-associated transcripts were considered during interpretation of differential gene expression analyses.

### Human lung tissue collection

Lung samples were collected from lung transplant recipients with CLAD undergoing retransplant or autopsy (after medical assistance in dying) between 2019 and 2024. All samples were collected following approved ethnical protocols and with documented informed consent and were processed within three hours of death or immediately following explantation. Prior to tissue processing, patients were provisionally classified as having a BOS or RAS phenotype based on pulmonary function testing and chest computed tomography findings. Cases with a clinically and radiologically clear BOS or RAS phenotype and without significant clinical complications were considered for scRNAseq. Practical considerations, including sample-processing capacity, personnel availability and sample quality influenced whether samples proceeded to scRNAseq. Following successful scRNAseq processing, the BOS or RAS phenotypes were retrospectively reviewed using available clinical, pulmonary function, radiographic and pathological data. Ultimately, 14 CLAD lung tissue (BOS = 7, RAS = 7) obtained from autopsy (n = 2) or retransplant cases (n = 12) successfully underwent scRNAseq.

### Lung tissue processing and scRNAseq

Lung tissue was processed into single-cell suspensions (20) and subjected to 3’ scRNAseq using the 10x Genomics (Pleasanton, CA, USA) platform, with downstream analysis performed using Cell Ranger and Seurat. Full analytical details are provided in the Online Supplement.

### Flow cytometry, cell sorting and functional assays

Single-cell suspensions from cryopreserved CLAD lung tissue and excess donor lung tissue were analyzed and sorted by spectral-enhanced fluorescence-activated cell sorting (FACS) to isolate macrophage subsets. Sorted macrophages were used for validation and in vitro functional assays. Full antibody panels, staining protocols, sorting strategies, validation procedures and in vitro assay methodologies are provided in the Online Supplement.

### Statistical analysis

Statistical analysis was performed with Prism 8 (Graphpad Software, San Diago, CA, USA). Group comparisons were conducted using Kruskal-Wallis test with post-hoc multiple comparisons as indicated. Data is presented as mean ± standard deviation, with statistical significance defined as p < 0.05.

### Study approval

Human biological samples were obtained through approved ethical protocols and with documented informed consent. This study was approved by the University Health Network Research Ethics Board (protocols 15-9531, 11-0509, 11-0170).

## Data availability

Single-cell RNA sequencing data generated in this study has been deposited in the Gene Expression Omnibus (GEO) under accession number TO BE DETERMINED. Analytical code used for data processing, analysis and figure generation is available at: https://github.com/Allen-Duong-uhn/CLAD-scRNAseq-macrophage.

## Impact Statement

By leveraging the largest single-cell analysis of chronic lung allograft dysfunction (CLAD) to date, this study identifies inflammatory and fibrotic macrophages enriched in CLAD. These populations were shared with other fibrotic lung diseases. . From the analysis, we derived surface marker-based strategies to isolate these macrophages and demonstrate their pathogenic functions using *in vitro* functional assays.

## Author Contributions

All authors contributed to Writing – Review/Editing.

**Allen Duong**: Conceptualization, Methodology, Software, Validation, Formal Analysis, Investigation, Writing (Original), Visualization.

**Sajad Moshkelgosha**: Conceptualization, Software.

**Aaron Wong**: Software, Resources.

**Ankita Burman**: Methodology, Resources

**Ke Fan Bei**: Methodology, Resources

**Rayoun Ramendra**: Methodology.

**Tsukasa Ishiwata**: Data Curation.

**Boris Hinz**: Methodology.

**Sonya MacParland**: Conceptualization, Resources.

**Mingyao Liu**: Resources.

**Stephen Juvet**: Conceptualization, Methodology, Resources, Supervision, Project administration, Funding acquisition.

**Tereza Martinu**: Conceptualization, Methodology, Resources, Supervision, Project administration, Funding acquisition.

## Funding/Financial support

Sanofi iAward (S.J, T.M)

Canadian Institutes of Health Research (CIHR) Project Grant (PJT 173343 and PJT 197913) (S.J, T.M)

CFF Grant (S.J, T.M)

Caron Thorburn Institute (S.J, T.M)

UHN Foundation (S.J., T.M.)

DiPoce Scholar Award (T.M.)

Truscott-Parkes Catalyst Fund in Idiopathic Pulmonary Fibrosis (S.J.)

## Acknowledgements

The authors thank the Toronto Lung Transplant Program Biobank team for their assistance with sample collection and retrieval, and the Toronto Lung Transplant Program Clinical Database team for their curation of clinical data. The authors also appreciate the hard work of the laboratory trainees and staff who have helped with lung tissue collection, processing and banking, including co-authors and also: Jamal Al-Refaee, Samuel Beber, Pavani Beesetty, Gregory Berra, Zoeen Carter, Nicole Chrysler, Tina Daignault, Jan Havlin, Mira Ishak, Mitsuaki Kawashima, Christina Lam, Liran Levy, Peter Lombardi, Goodness Madu, Olivia Mekhael, Ei Miyamoto, Gafoor Puthiyaveetil, Sumiha Ramendra, Benjamin Renaud-Picard, Nadia Sachewsky, David Sebben, Julie Semenchuk, Yamato Suzuki, Akihiro Takahagi, Daniel Vosoughi, Tatsuaki Watanabe, Reese Whittaker, Matthew White, Kevin Zhang, Wenshan Zhong, and Nancy Zhao. Library creation and next-generation sequencing were performed by the Princess Margaret Genomic Centre (RRID: SCR_027524). Spectral-enhanced fluorescence-activated cell sorting was performed by the University of Toronto Flow Cytometry Facility (RRID: SCR_027612). The authors thank the Princess Margaret Flow Cytometry Facility (RRID: SCR_027537) for the use of their analytical spectral flow cytometer and the University Health Network’s Advanced Optical Microscopy Facility (RRID: SCR_027535) for the use of their microscopes.

## Abbreviations

AMΦ: alveolar macrophage
BAL: bronchoalveolar lavage
BOS: bronchiolitis obliterans syndrome
CLAD: chronic lung allograft dysfunction
FACS: fluorescence-activated cell sorting
IPF: idiopathic pulmonary fibrosis
IFNγ: interferon-γ
IL: interleukin
IMΦ: interstitial macrophage
ISG: interferon-stimulated genes
LILR: leukocyte immunoglobulin-like receptor
RAS: restrictive allograft syndrome
RT-qPCR: reverse transcription-quantitative polymerase chain reaction
scRNAseq: single-cell RNA sequencing
SPP1: secreted phosphoprotein 1
TGF-β: transforming growth factor-beta

