## Supplemental Material for "Identification and characterization of inflammatory LILR and fibrotic SPP1 macrophages in chronic lung allograft dysfunction"

### ONLINE DATA SUPPLEMENT

### SUPPLEMENTARY METHODS

*Lung tissue dissociation*

Explanted CLAD lungs were collected and dissected, obtaining two lung tissue samples (1 x 4 x 4 cm) from a depth of at least 4 cm from the pleural surface. Lung tissue samples were processed as previously described (1). Briefly, lung tissue was minced and transferred to gentleMACS C Tubes (Miltenyi Biotec, Bergisch Gladbach, Germany) containing 5 ml Hanks Buffered Salt Solution supplemented with 5% vol/vol heat inactivated fetal bovine serum (HI-FBS) (Gibco, Waltham, MA, USA), 10 mM 4-(2-hydroxyethyl)-1-piperazineethansulfonic acid (HEPES), 800 µg/ml Collagenase A (Millipore Sigma, Burlington MA, USA) and 160 µg/ml DNase I (Millipore Sigma). Samples underwent dissociation using a gentleMACS Octo Dissociator (Miltenyi Biotec) following a custom program described previously (1). After dissociation, samples were filtered through a 70 µm cell strainer and resuspended in erythrocyte lysis solution (155 mM $NH_4Cl$,10 mM $KHCO_3$, 68.5 µM ethylenediaminetetraacetic acid. (EDTA)) for 5 min. Cells were then washed with flow cytometry staining buffer (FACS buffer; phosphate-buffered saline (PBS; Wisent Bioproducts, Saint-Jean Baptiste, Canada) with 2% vol/vol HI-FBS and 2 mM EDTA) and quantified through manual cell counting using a hemocytometer with trypan blue exclusion. Final single cell suspensions were resuspended in PBS with 0.04% bovine serum albumen (BSA; Sigma Aldrich).

*Capture, library construction and sequencing*

A total of 11 µl single cell suspension at a concentration of 1000 cells/µl and 40 µl of barcoded gel beads were loaded onto a Chromium Chip (10x Genomics, Pleasanton, CA, USA) to generate a single-cell gel bead emulsion, targeting the recovery of 10000 cells. Single cell capture and downstream library construction were performed using 10x Genomics 3' expression v3.1 chemistry. Full length cDNA and cell-barcode identifiers were amplified through PCR and sequencing libraries prepared. The constructed libraries were frozen and later sequenced on the Illumina® NovaSeq platform (Illumina, San Diego, CA, United States).

*scRNAseq data alignment and quality control*

Cell Ranger (v7.2.0, 10x Genomics) was used to perform sample demultiplexing, barcode processing and single cell 3' unique molecular identifier (UMI) counting with human GRCh38.2020 as the reference genome. Splicing-aware aligner STAR was used for FASTQ alignment. Cell barcodes were determined based on the distribution of UMI counts automatically. Each individual sample was processed through Seurat (2) (v5.3.0) package in R and underwent quality control, using SoupX (3) to remove ambient RNA, removing genes encoding ribosomal proteins (genes starting with RPL or RPS), using DoubletFinder (4) to remove doublets, and filtering cells with high mitochondrial content (greater than 15%) and small or excessively high library sizes (gene number lower than 200 or higher than 5000, UMI counts lower than 1000 or higher than 30000). After filtration, the remaining cells were subjected to subsequent analysis.

*Cell annotation*

Cells were initially annotated using the SingleR package (5) and then manually reviewed using markers identified in the pulmonary biology literature (6) (Figure S1A). Annotations were added to Seurat object as meta-data.

*scRNAseq integration and subsetting*

The 7 BOS, 7 RAS and 6 controls datasets were merged and integrated using Seurat sctransform v2 workflow (7), and preprocessing performed as previously stated. All clusters with cell annotation “macrophage” or “monocyte” were subsetted and reclustered, using a resolution of 1.0, as determined through Clustree package (8) to avoid over-clustering.

Differential gene expression analysis

Seurat FindAllMarkers function using Wilcoxon rank sum test was used to identify the top ten differential gene expression by average log2 fold change for each macrophage cluster, with the following cut-offs: min.pct = 0.25, logfc.threshold = 0.25. To identify differential gene expression between CLAD versus control and BOS versus RAS, model-based analysis of single-cell transcriptomics (MAST) analysis (9) was performed in Seurat v5.

*Cell-cell interaction analysis*

To infer intercellular communication, R package CellChat (10) (v2.1.0) was used on normalized lung scRNAseq data. Statistically significant interactions ($p < 0.05$) were identified and quantified. The top outgoing signaling pathways originating from macrophages were identified using “netAnalysis_signalingRole_heatmap”. The inferred cell targets were visualized as a circle plot through function “netVisual_aggregate” and the specific contributions of each ligand-receptor pair for the top outgoing signaling pathways were identified using function “netAnalysis_contribution”.

*Pathway analysis*

Gene ontology (GO) and Gene Set Enrichment Analysis (GSEA) was performed using clusterProfiler package (11) (v4.14.6) in R to identify significantly enriched ($p < 0.05$) biological processes, molecular functions, and pathways among differentially expressed genes within each macrophage subset.

*Gene regulatory network (GRN) analysis*

To identify enriched GRNs and active transcription factors across macrophages clusters, SCENIC (12) (v1.3.1) implemented in R was used. The SCENIC pipeline under default parameters was applied to a macrophage Seurat object downsampled to 500 cells per annotated cluster to identify transcription factor-target interactions and estimate regulon activity.

*Fluorescence activated cell sorting (FACS)*

R package cerebroApp was used to identify transcripts encoding cell-surface marker proteins, which were used to sort specific macrophage populations. SDC2 was identified to sort SPP1 macrophages, and LILRB2 was used to sort LILR macrophages. CLAD lung single cell suspensions were stained using a validation sorting panel (antibodies detailed in Table S2). Flow cytometry staining protocol was performed as previously published (1). A Sony SH800 cell sorter (Sony Biotechnology, San Jose, CA, USA) was used to sort SPP1 and LILR macrophages for validation purposes.

*Sorting validation*

After sorting SPP1 and LILR macrophages, the populations were subsequently stained with an analytical flow cytometry panel, detailed in Table S2. The analytical flow cytometry panel includes the antibodies in the sorting panel but is expanded to include intracellular anti-SPP1- and anti-AIF1 antibodies. Staining procedure is as stated above, with the addition of intracellular staining using the BD Cytofix/Cytoperm™ Fixation/Permeabilization Kit (BD Bioscience, Franklin Lakes, NJ, USA) following manufacturer's instructions. Flow cytometry was run on a ID7000 spectral flow cytometer (Sony Biotechnology).

To further validate that the sorted cells were the identified scRNAseq clusters, reverse transcription-quantitative PCR (RT-qPCR) was performed, targeting transcripts identified by scRNAseq DEG analysis. Sorted cells underwent RNA extraction using TRIzol™

reagent (Thermo Fisher Scientific, Waltham, MA, USA) and purification through RNeasy Minikit (Qiagen, Hilden, Germany). Purified RNA was converted to complimentary DNA (cDNA) using iScript™ cDNA synthesis kit (BioRAD, Hercules, CA, USA) and qPCR was performed using SsoAdvanced Universal SYBR Green Supermix (BioRAD) with primers targeting: *ACTB*, *AIF1*, *LILRB2*, *MMP9*, *SDC2* (specific primer sequences reported in Table S3). Resultant RT-qPCR data was analyzed using CFX Maestro Software for Real-Time PCR (BioRAD) and normalized to *ACTB* expression.

*Cell sorting for culturing*

Thawed single cell suspensions of CLAD (n = 12, 6 BOS, 6 RAS) and non-CLAD donor excess lung samples (n = 4) were stained with sorting panel (antibodies detailed in Table S2) using the same staining protocol. Spectral-enhanced FACS was performed using a FACSymphony™ S6 (BD Bioscience) and LILR+, SPP1+ and LILR-SPP1- (Other) macrophages were isolated. Following isolation, macrophages were cultured at 1 x $10^5$ cells/ml in multiple wells of an untreated 96-well plates (Corning Inc., Corning, NY, USA) as technical replicates with Dulbecco's Modified Eagle Medium (DMEM; Thermo Fisher) supplemented with 10% volume/volume (v/v) fetal bovine serum (FBS; Thermo Fisher), 100 U/ml penicillin, 100 µg/ml streptomycin, 0.1mM minimum essential medium non-essential amino acids (gibco®, Thermo Fisher), 2mM GlutaMAX (gibco®, Thermo Fisher) ,1mM sodium pyruvate (gibco) and 0.05mM 2-mercaptoethanol. Macrophages were cultured for 24 hours, after which conditioned supernatants were harvested, pooled across technical replicates, and cryopreserved in -20 °C. Fresh culture media was then added to the cells.

*Phagocytosis assay*

Following supernatant harvest, the macrophages phagocytotic ability was assessed using pHrodo™ Red *E. coli* BioParticles™ (Thermo Fisher) at the manufacturer suggested concentration. Immediately following addition of BioParticles™, the macrophages were imaged using an Axio Observer widefield microscope (Zeiss, Oberkochen, Germany) equipped with Live Cell Instrument Chamlide stage-top incubator, using 10x phase contrast objective lense and Cy3 fluorescence filter set. Images were taken every 15 mins for 2 hours. Images were processed in ZEN Blue (Zeiss).

*In vitro fibrosis assays*

The pulmonary fibroblast cell line MRC-5 (American Type Culture Collection (ATCC), Manassas, VA, USA) was used for all *in vitro* fibroblast assays. Low passage MRC-5 cells were cultured in untreated 24-well plates with complete DMEM with 10% v/v of macrophage culture supernatant (LILR macrophage, SPP1 macrophage, Other macrophage and saline control) until 90% confluent. MRC-5 cells cultured with 2 ng/ml TGFβ (Peprotech, Cranbury, NJ, USA) for 3 days were used as positive controls. A scratch migration assay was performed, using a 200 μl pipette tip to create a gap in the fibroblast monolayer. The plate was placed in an Axio Observer widefield microscope (Zeiss) with incubator system, and images of the gap were obtained every 30 mins for 24 hours. Gap width was measured at nine locations within each well and averaged.

Fibroblast migration was expressed as the percentage reduction in gap width relative to baseline (0 hours).

To measure fibroblast contraction following migration imaging, the MRC-5 cells were detached from the well using 0.25% Trypsin in EDTA (gibco®, Thermo Fisher), and 175,000 cells were mixed with 100 µl of 6 mg/ml bovine dermal type I collagen (Advanced BioMatrix, San Diago, CA, USA). Collagen-fibroblast mixtures were then neutralized with 0.1M NaOH to pH 7 and pipetted into 35 mm petri dishes (Corning) and allowed to solidify (13). After the gel solidified, 3.5 ml of DMEM was added to prevent drying, and the dishes were placed into the incubator (37 °C 5% $CO_2$) for 5 days. After incubation, the attached gels were imaged, then and detached from the petri dish, and imaged again every 30 min for 90 min.

*Endothelial cell assay*

Low-passage HULEC-5a (ATCC), microvascular pulmonary endothelial cell line was cultured in untreated 24-well plates with complete DMEM with 10% v/v of macrophage culture supernatant (LILR macrophage, SPP1 macrophage, Other macrophage and saline control). HULEC-5a cells cultured with TNF-α (10 ng/ml) or IFN-γ (250 ng/ml) were used as positive controls. Following 3 days of culture, HULEC-5a cells were detached with 0.25% Trypsin-EDTA (Gibco®, Thermo Fisher), and underwent flow cytometry staining for activation markers (Table S4). Analytical flow cytometry was performed on the Sony ID7000 analytical spectral flow cytometer (Sony Biotechnologies) and analyzed via FlowJo X (BD Bioscience, San Jose, CA, USA).

*T cell chemotaxis assay*

PBMCs were isolated from blood provided by volunteers using the SepMate™ PBMC isolation tubes with Lymphoprep™ density gradient medium (StemCell Technologies, Vancouver, BC, Canada), following manufacturer's protocol. T cells were subsequently isolated from PBMCs using EasySep™ human T cell magnetic isolation kit (StemCell Technologies). T cells were cultured in non-treated 24-well plates with Roswell Park Memorial Institute 1640 (RPMI; gibco®, Thermo Fisher) supplemented with 10% v/v FBS (Thermo Fisher), 100 U/ml penicillin, 100 µg/ml streptomycin, 0.1mM minimum essential medium non-essential amino acids (gibco), 2mM GlutaMAX® (gibco) ,1mM sodium pyruvate (gibco) and 0.05mM 2-mercaptoethanol, and activated with Dynabeads™ Human T-Activator CD3/CD28 (Thermo Fisher) and recombinant human IL-2 (100 IU/ml). After 4 days of culture, the Dynabeads™ were removed via magnetic field and T cells resuspended in cRPMI.

T cell chemotaxis was measured using Transwells® inserts (Corning) with 8µm pore size for 24-well plates. cRPMI with 10% v/v of macrophage culture supernatant (LILR, SPP1, Other macrophages and saline control) or 1000ng/ml CXCL10 were placed at the bottom chamber. 250,000 activated T cells were placed at the top chamber and allowed to incubate overnight at standard cultures.

*Cytokine multiplex assay*

To measure cytokines generated by the cultured macrophage supernatants, supernatants were profiled using the 96-Plex Discovery assay (Human Cytokine/Chemokine Array 96-plex Panel; Cat no. HD96; Eve Technologies, Calgary, AB, Canda). The initial 96 analytes were subsequently filtered to exclude those for which more than 50% of samples below the lower limit of quantification. The following analytes met inclusion criteria and were retained for analysis: Epidermal growth factor (EGF), eotaxin, FMS-like tyrosine kinase 3 ligand (FLT-3L), granulocyte colony-stimulating factor (G-CSF), granulocyte–macrophage colony-stimulating factor (GM-CSF), growth-related oncogene-α (GROα/CXCL1), IL-1α, IL-1β, IL-1 receptor antagonist (IL-1RA), IL-2, IL-3, IL-6, IL-8, IL-10, IL-12p40, IL-12p70, IL-13, IL-18, IL-22, interferon gamma–induced protein 10 (IP-10/CXCL10), monocyte chemoattractant protein-1 (MCP-1/CCL2), MCP-3 (CCL7), monokine induced by interferon-γ (MIG/CXCL9), macrophage inflammatory protein-1α (MIP-1α/CCL3), MIP-1β (CCL4), transforming growth factor-α (TGF-α), tumor necrosis factor-α (TNF-α), TNF-β, vascular endothelial growth factor-A (VEGF-A), a proliferation-inducing ligand (APRIL), CXCL16, epithelial neutrophil–activating peptide 78 (ENA-78/CXCL5), eotaxin-2 (CCL24), eotaxin-3 (CCL26), granulocyte chemotactic protein-2 (GCP-2/CXCL6), high mobility group box 1 (HMGB1), I-309 (CCL1), interferon-ω (IFN-ω), IL-16, IL-20, IL-21, IL-23, lymphotactin (XCL1), MCP-2 (CCL8), MIP-3α (CCL20), myeloid progenitor inhibitory factor-1 (MPIF-1/CCL23), stem cell factor (SCF), thymus and activation-regulated chemokine (TARC/CCL17), and thymic stromal lymphopoietin (TSLP). Samples falling beyond the lower range of quantification and unsuitable for extrapolation were assigned a value of the lowest with fluorescence indices fallout out of range of the standard curve were assigned a value of one-fifth of the lowest observed concentration for the respective analyte.

### SUPPLEMENTARY FIGURES

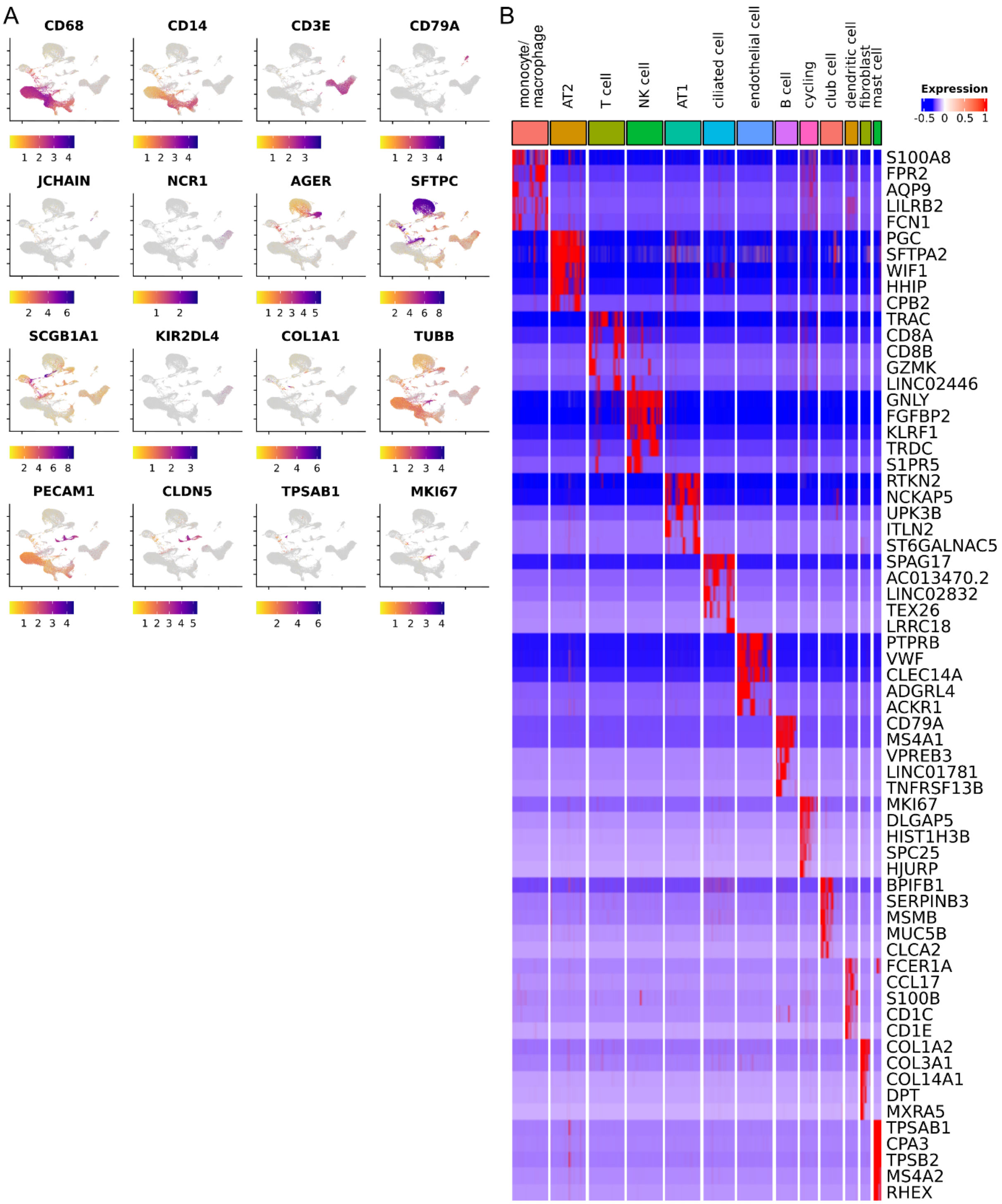


**Supplementary Figure 1: Canonical marker gene expression used for lung cell type annotation of single-cell RNA sequencing data from 14 CLAD lungs (BOS = 7, RAS**

**= 7) and 6 control lungs**. A) Feature plots showing relative expression of canonical marker genes: CD68, CD14, CD3E, CD79A, JCHAIN, NCR1, AGER, SFTPC, SCGB1A1, KIR2DL4, COL1A1, TUBB, PECAM1, CLDN5, TSPAB1, MKI67. B) Heatmap depicting the top five differentially expressed genes from each annotated lung cell population.

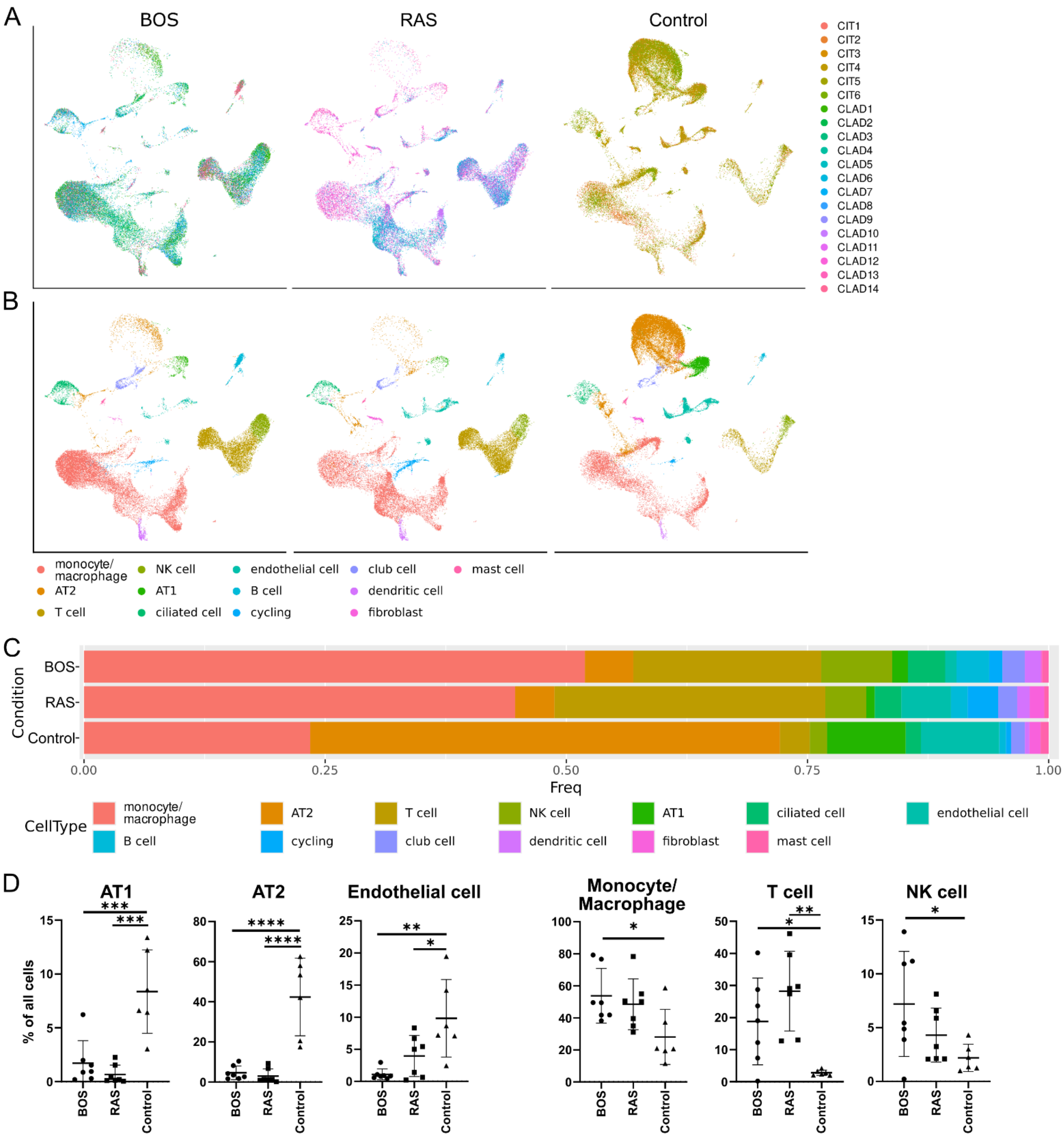


**Supplementary Figure 2: Comparative analysis of major pulmonary cell populations from single-cell RNA sequencing data across BOS, RAS and Control**

**lungs.** A and B) UMAP visualization of pulmonary cells split by BOS, RAS or Control lung, grouped by A) Biological replicate and B) annotated cell type. C) Stacked bar plot showing cellular composition within BOS, RAS and Control lungs. D) Proportional representation of type I alveolar cell (AT1), type 2 alveolar cell (AT2), endothelial cell, monocyte/macrophage, T cells, and NK cells across individual lung samples. Group comparisons were performed using a Kruskal-Wallis test, with post hoc multiple comparisons performed and reported as follows: * $P < 0.05$, ** $P < 0.01$, *** $P < 0.001$, **** $P < 0.0001$.

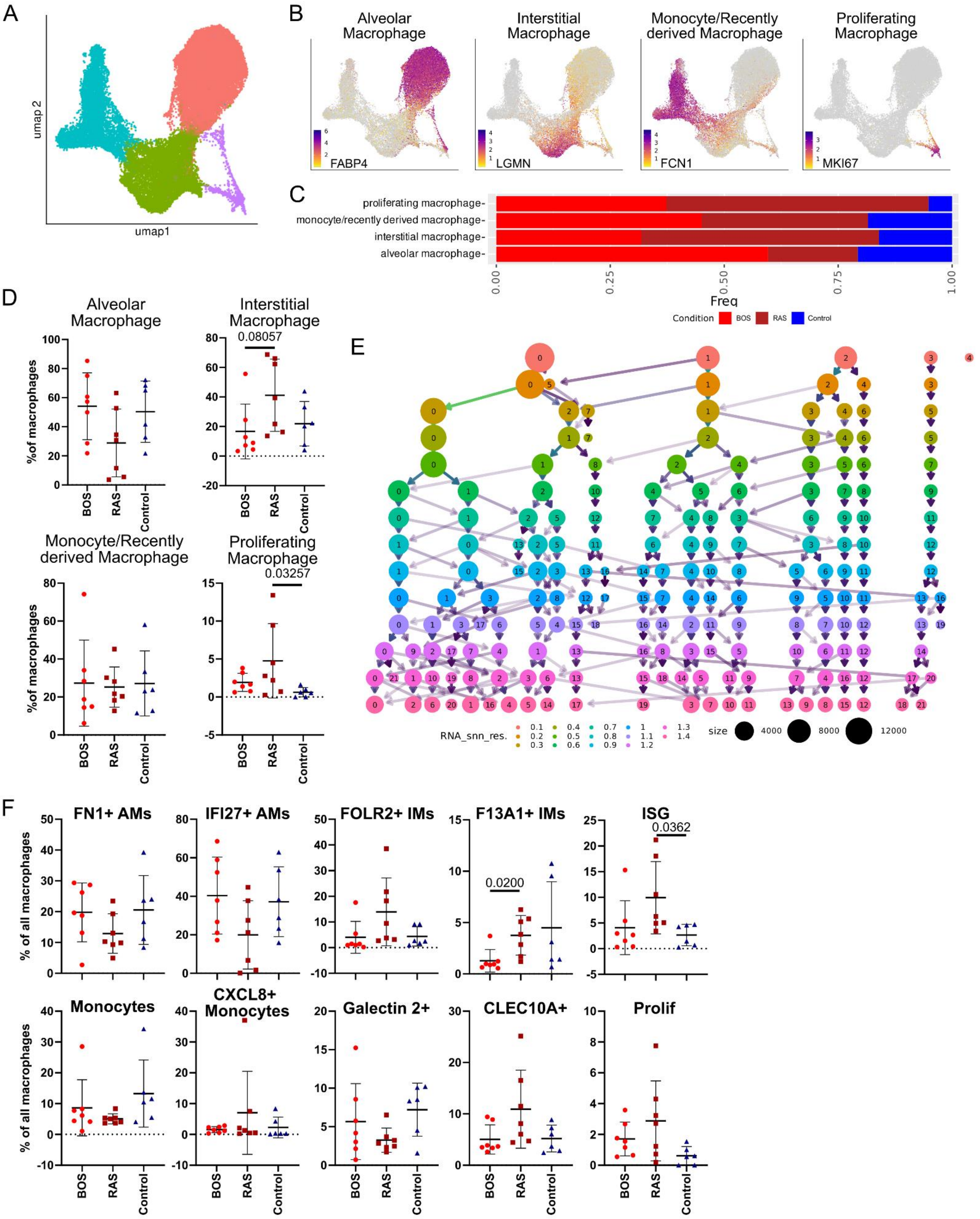
A
umap2
umap1
B
Alveolar Macrophage
FABP4
Interstitial Macrophage
LGMN
Monocyte/Recently derived Macrophage
FCN1
Proliferating Macrophage
MKI67
C
proliferating macrophage-
monocyte/recently derived macrophage-
interstitial macrophage-
alveolar macrophage-
0.00
0.25
0.50
0.75
1.00
Freq
Condition BOS RAS Control
D
Alveolar Macrophage
Interstitial Macrophage
0.08057
Monocyte/Recently derived Macrophage
Proliferating Macrophage
0.03257
%of macrophages
BOS
RAS
Control
E
RNA_snn_res.
0.1 0.2 0.3 0.4 0.5 0.6 0.7 0.8 0.9 1 1.1 1.2 1.3 1.4
size 4000 8000 12000
F
FN1+ AMs
IFI27+ AMs
FOLR2+ IMs
F13A1+ IMs
0.0200
ISG
0.0362
Monocytes
CXCL8+ Monocytes
Galectin 2+
CLEC10A+
Prolif
% of all macrophages
BOS
RAS
Control

**Supplementary Figure 3: Hierarchical clustering and proportional distribution of monocyte and macrophage subsets in CLAD lung tissue analyzed by single-cell RNA sequencing.** A) UMAP visualization showing low-resolution clustering of monocytes and macrophages, identifying four major annotated myeloid populations. B) Feature plots showing canonical marker gene expression of *FABP4*, *LGMN*, *FCN1* and *MKI67*, supporting the distinction between alveolar macrophages, interstitial macrophages, monocyte/recently monocyte-derived macrophages, and proliferating macrophages. C) Stacked bar plot showing the proportional distribution of the four major monocyte/macrophage groups, expressed as the percentage of total monocytes/macrophages within each group. D) Proportional representation of the four major monocyte/macrophage groups across BOS, RAS, and control samples, shown as the percentage of each group contributed by each phenotype. E) Clustree analysis illustrating hierarchical clustering of monocytes and macrophages across increasing resolution parameters (0.1–1.4), supporting the use of 1.0 resolution. F) Proportional representation of the ten higher-resolution macrophage subsets across BOS, RAS, and control samples.

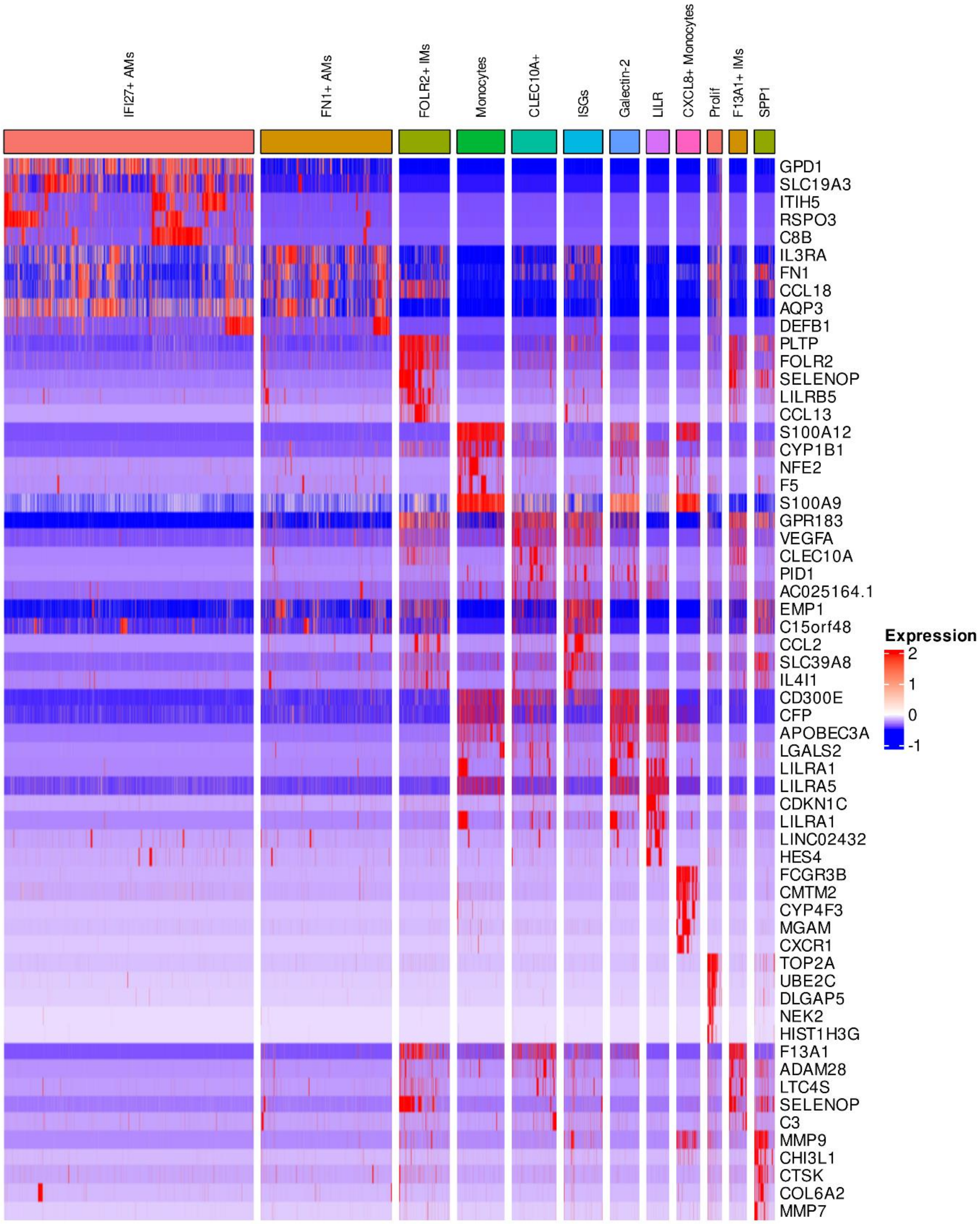


**Supplementary Figure 4: Heatmap depicting the top five differentially expressed genes from each annotated macrophage subset from single-cell RNA sequencing data from CLAD and control lungs.**

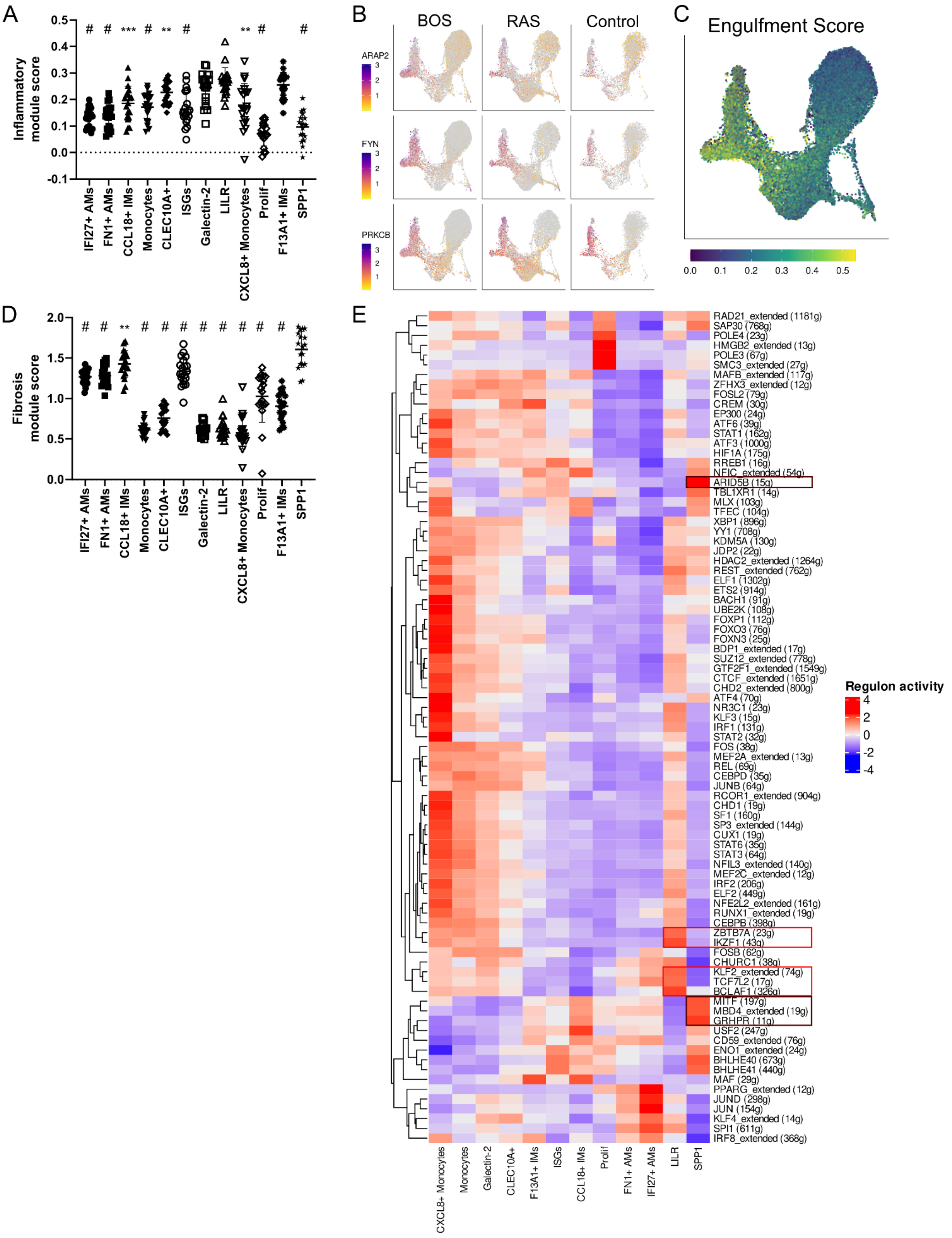
A
Inflammatory module score
# # *** # ** # ** # #
IFI27+ AMs
FN1+ AMs
CCL18+ IMs
Monocytes
CLEC10A+
ISGs
Galectin-2
LILR
CXCL8+ Monocytes
Prolif
F13A1+ IMs
SPP1
B
BOS
RAS
Control
ARAP2
FYN
PRKCB
C
Engulfment Score
D
Fibrosis module score
# # ** # # # # # # # # #
E
Regulon activity
RAD21_extended (1181g)
SAP30 (768g)
POLE4 (23g)
HMGB2_extended (13g)
POLE3 (67g)
SMC3_extended (27g)
MAFB_extended (117g)
ZFHX3_extended (12g)
FOSL2 (79g)
CREM (30g)
EP300 (24g)
ATF6 (39g)
STAT1 (162g)
ATF3 (1000g)
HIF1A (175g)
RREB1 (16g)
NFIC_extended (54g)
ARID5B (15g)
TBL1XR1 (14g)
MLX (103g)
TFEC (104g)
XBP1 (896g)
YY1 (708g)
KDM5A (130g)
JDP2 (22g)
HDAC2_extended (1264g)
REST_extended (762g)
ELF1 (1302g)
ETS2 (914g)
BACH1 (91g)
UBE2K (108g)
FOXP1 (112g)
FOXO3 (76g)
FOXN3 (25g)
BDP1_extended (17g)
SUZ12_extended (778g)
GTF2F1_extended (1549g)
CTCF_extended (1651g)
CHD2_extended (800g)
ATF4 (70g)
NR3C1 (23g)
KLF3 (15g)
IRF1 (131g)
STAT2 (32g)
FOS (38g)
MEF2A_extended (13g)
REL (69g)
CEBPD (35g)
JUNB (64g)
RCOR1_extended (904g)
CHD1 (19g)
SF1 (160g)
SP3_extended (144g)
CUX1 (19g)
STAT6 (35g)
STAT3 (64g)
NFIL3_extended (140g)
MEF2C_extended (12g)
IRF2 (206g)
ELF2 (449g)
NFE2L2_extended (161g)
RUNX1_extended (19g)
CEBPB (398g)
ZBTB7A (23g)
IKZF1 (43g)
FOSB (62g)
CHURC1 (38g)
KLF2_extended (74g)
TCF7L2 (17g)
BCLAF1 (326g)
MITF (197g)
MBD4_extended (19g)
GRHPR (11g)
USF2 (247g)
CD59_extended (76g)
ENO1_extended (24g)
BHLHE40 (673g)
BHLHE41 (440g)
MAF (29g)
PPARG_extended (12g)
JUND (298g)
JUN (154g)
KLF4_extended (14g)
SPI1 (611g)
IRF8_extended (368g)
CXCL8+ Monocytes
Monocytes
Galectin-2
CLEC10A+
F13A1+ IMs
ISGs
CCL18+ IMs
Prolif
FN1+ AMs
IFI27+ AMs
LILR
SPP1

**Supplementary Figure 5: Transcriptional activity, phagocytic capacity, and regulatory programs across macrophage subsets from single-cell RNA sequencing data from CLAD and control lungs.** A) Dot plot showing inflammatory gene module scores across macrophage subsets. Statistical comparisons were performed using a Kruskal-Wallis test comparing each subset to LILR macrophages. Significant differences are indicated by symbols ** $P < 0.01$, *** $P < 0.001$, # $P < 0.0001$. B) Feature plots showing relative expression of representative phagocytosis-related genes: ARAP2, FYN and PRKCB. C) UMAP visualization of enrichment scores for engulfment gene programs. D) Dot plot showing fibrotic gene module scores across macrophage subsets. Statistical comparisons were performed using a Kruskal-Wallis test comparing each subset to SPP1 macrophages. Significant differences are indicated by symbols ** $P < 0.01$, # $P < 0.0001$. E) Heatmap showing top regulon activity across macrophage subsets as inferred by SCENIC analysis.

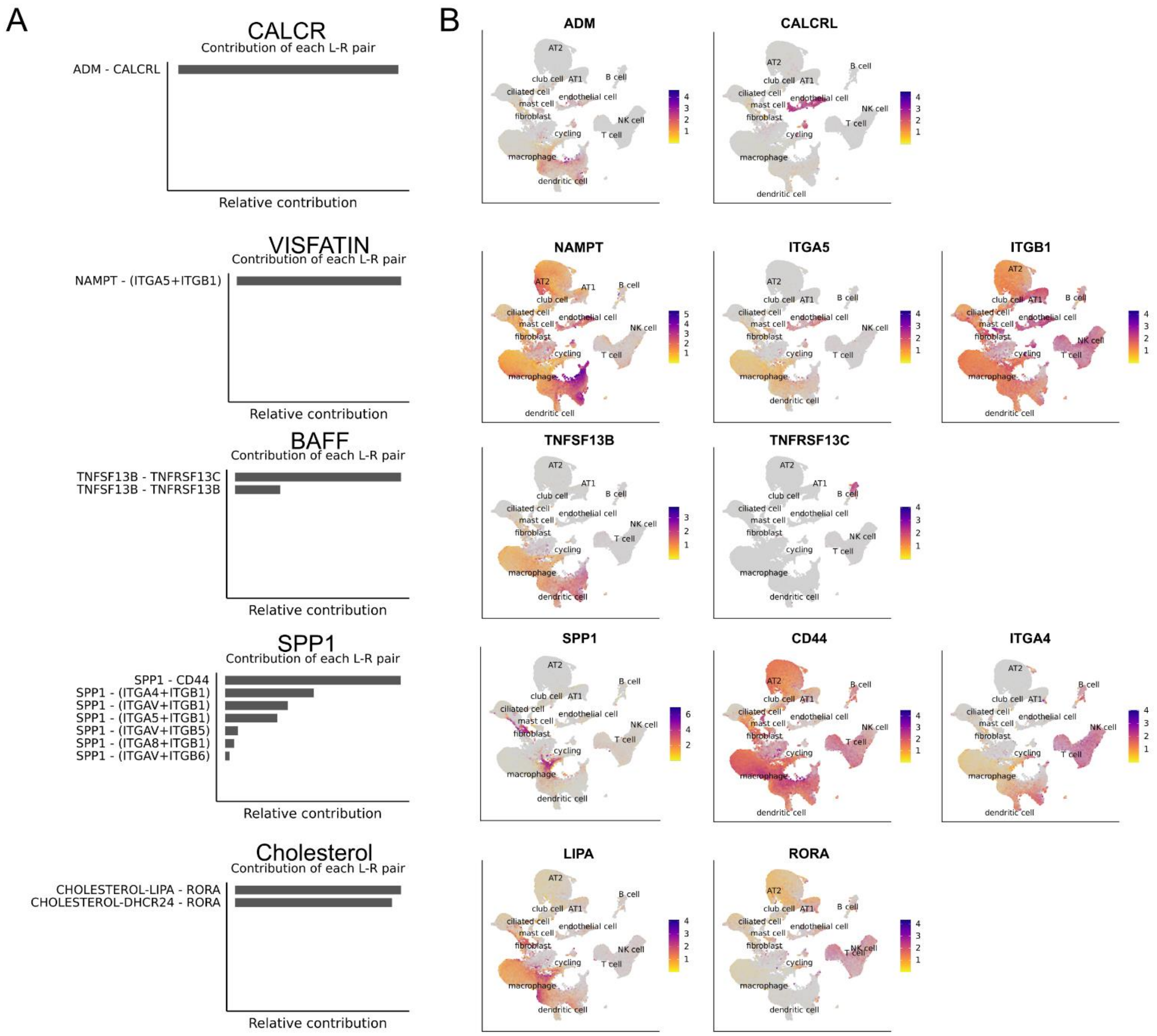


**Supplementary Figure 6: Expression of ligand–receptor gene pairs underlying CellChat-inferred signaling pathways in LILR and SPP1 macrophages from single-cell RNA sequencing data from CLAD and control lungs.** A) Relative contribution of ligand-receptor gene pairs from the CALCR, VISFATIN, BAFF, SPP1 and Cholesterol signaling pathways. B) Feature plots showing the expression of the corresponding ligand and receptor pairs identified in panel A.

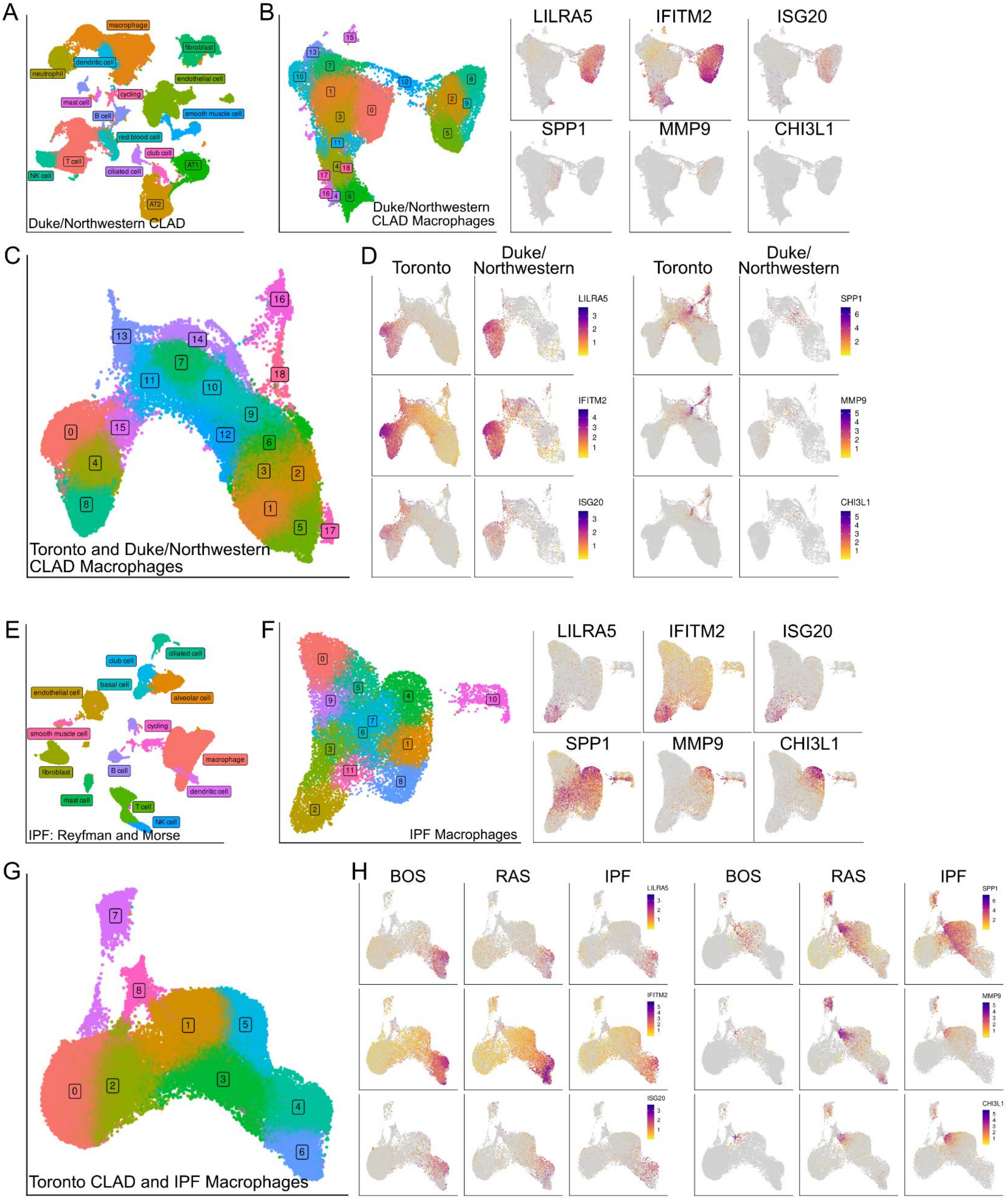

A
macrophage
fibroblast
dendritic cell
neutrophil
endothelial cell
mast cell
cycling
B cell
smooth muscle cell
red blood cell
club cell
T cell
AT1
NK cell
ciliated cell
AT2
Duke/Northwestern CLAD
B
Duke/Northwestern
CLAD Macrophages
LILRA5
IFITM2
ISG20
SPP1
MMP9
CHI3L1
C
Toronto and Duke/Northwestern
CLAD Macrophages
D
Toronto
Duke/
Northwestern
E
club cell
ciliated cell
basal cell
endothelial cell
alveolar cell
cycling
smooth muscle cell
macrophage
fibroblast
B cell
dendritic cell
mast cell
T cell
NK cell
IPF: Reyfman and Morse
F
IPF Macrophages
G
Toronto CLAD and IPF Macrophages
H
BOS
RAS
IPF

**Supplementary Figure 7: Comparison of inflammatory and fibrotic macrophage programs across idiopathic pulmonary fibrosis (IPF) and independent CLAD single-cell RNA sequencing datasets.** A) Annotated UMAP visualization of lung cell populations from IPF lung tissue. B) UMAP visualization of macrophages isolated from IPF lungs (left), with feature plots showing expression of selected pro-inflammatory (LILRA5, IFITM2, ISG20) and pro-fibrotic (SPP1, MMP9, CHI3L1) marker genes used to identify LILR and SPP1 macrophage populations (right). C) UMAP visualization of integrated clustering of macrophages from Toronto CLAD and IPF lung datasets. D) Feature plots of the same marker genes shown in panel B, visualized in the integrated Toronto CLAD–IPF macrophage dataset and stratified by BOS, RAS, and IPF samples. E) Annotated UMAP visualization of lung cell populations from publicly available CLAD lung scRNA-seq datasets (Duke and Northwestern cohorts). F) UMAP visualization of macrophages isolated from IPF lungs (left), with feature plots showing expression of selected pro-inflammatory and pro-fibrotic marker genes (right). G) UMAP visualization of integrated clustering of macrophages from Toronto and Duke/Northwestern CLAD datasets. H) Feature plots of the same marker genes shown in panel B and F, visualized in the integrated CLAD datasets and stratified by cohort.

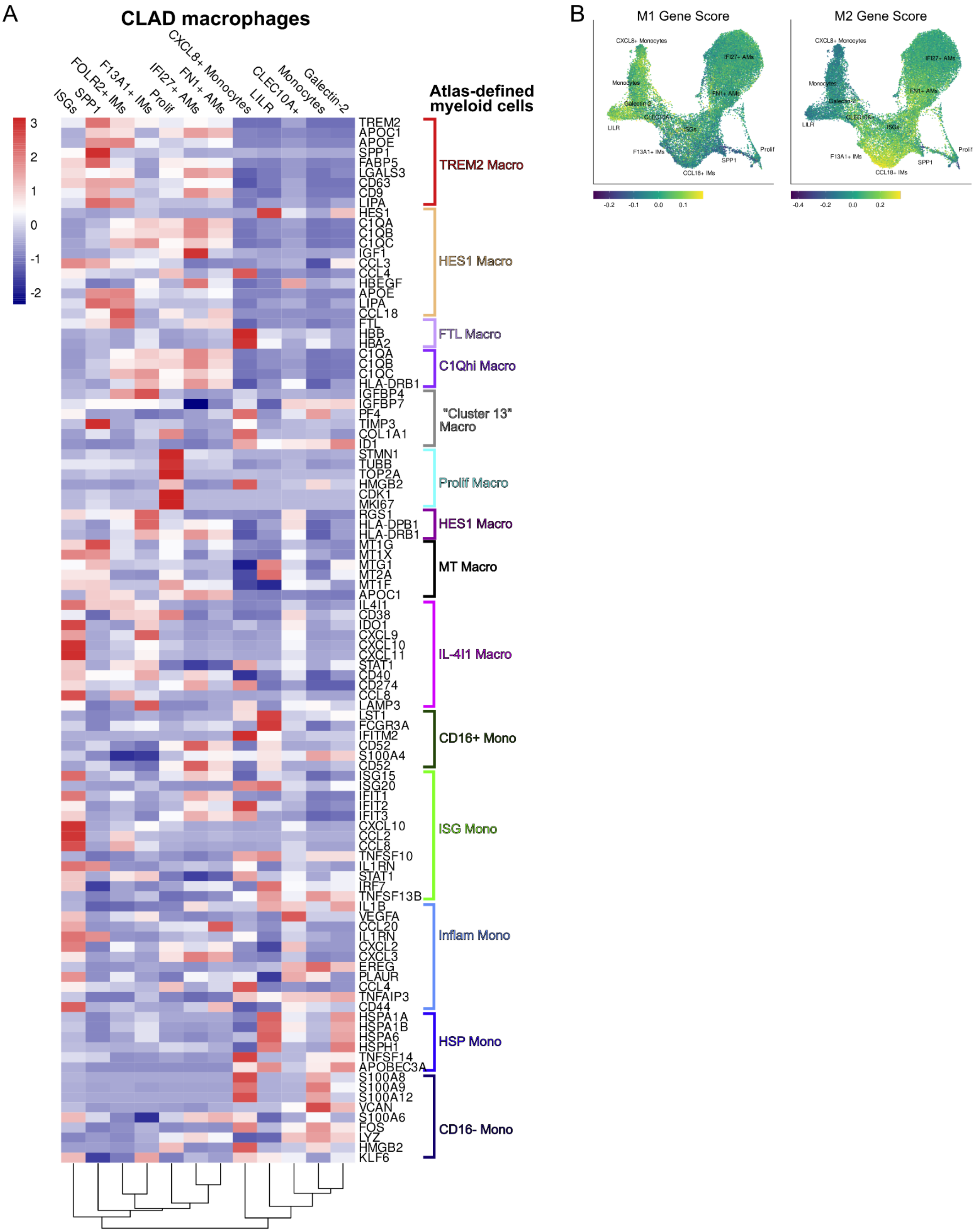
A
CLAD macrophages
ISGs
SPP1
FOLR2+ IMs
F13A1+ IMs
Prolif
IFI27+ AMs
FN1+ AMs
CXCL8+ Monocytes
LILR
CLEC10A+
Monocytes
Galectin-2
Atlas-defined myeloid cells
3
2
1
0
-1
-2
TREM2 APOC1 APOE SPP1 FABP5 LGALS3 CD63 CD9 LIPA
TREM2 Macro
HES1 C1QA C1QB C1QC IGF1 CCL3 CCL4 HBEGF APOE LIPA CCL18
HES1 Macro
FTL HBB HBA2
FTL Macro
C1QA C1QB C1QC HLA-DRB1
C1Qhi Macro
IGFBP4 IGFBP7 PF4 TIMP3 COL1A1 ID1
"Cluster 13" Macro
STMN1 TUBB TOP2A HMGB2 CDK1 MKI67
Prolif Macro
RGS1 HLA-DPB1 HLA-DRB1
HES1 Macro
MT1G MT1X MTG1 MT2A MT1F APOC1
MT Macro
IL4I1 CD38 IDO1 CXCL9 CXCL10 CXCL11 STAT1 CD40 CD274 CCL8 LAMP3
IL-4I1 Macro
LST1 FCGR3A IFITM2 CD52 S100A4 CD52
CD16+ Mono
ISG15 ISG20 IFIT1 IFIT2 IFIT3 CXCL10 CCL2 CCL8 TNFSF10 IL1RN STAT1 IRF7 TNFSF13B
ISG Mono
IL1B VEGFA CCL20 IL1RN CXCL2 CXCL3 EREG PLAUR CCL4 TNFAIP3 CD44
Inflam Mono
HSPA1A HSPA1B HSPA6 HSPH1 TNFSF14 APOBEC3A
HSP Mono
S100A8 S100A9 S100A12 VCAN S100A6 FOS LYZ HMGB2 KLF6
CD16- Mono
B
M1 Gene Score
M2 Gene Score
CXCL8+ Monocytes
IFI27+ AMs
FN1+ AMs
Monocytes
Galectin-2
CLEC10A+
ISGs
LILR
F13A1+ IMs
SPP1
CCL18+ IMs
Prolif
-0.2
-0.1
0.0
0.1
-0.4
-0.2
0.0
0.2

**Supplementary Figure 8: Cross-tissue macrophage signatures in single-cell RNA sequencing data map LILR and SPP1 subsets beyond classical M1/M2 states.** A) Heatmap showing average expression of selected macrophage marker genes derived from cross-tissue scRNAseq framework described by Mulder et al. Marker genes are grouped according to macrophage/monocyte categories defined in the Mulder atlas, and columns represent macrophage subsets. Gene expression is averaged across cells within each macrophage subset. B) UMAP visualization of enrichment scores for M1 and M2 gene modules.

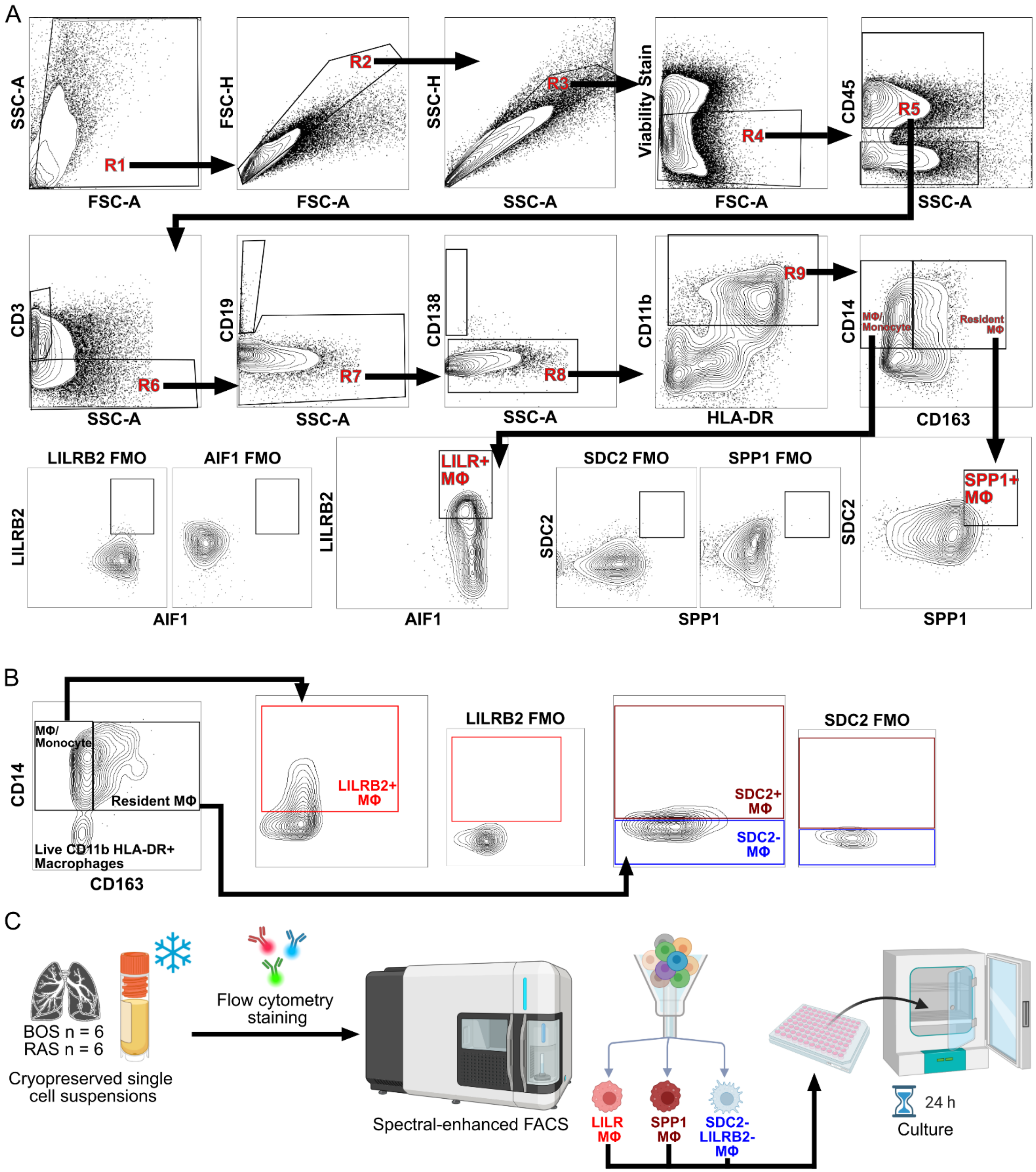


**Supplementary Figure 9: Flow cytometry gating, sorting strategies, and experimental workflow for isolation of LILR and SPP1 macrophages (MΦ).** A)

Analytical flow cytometry gating strategy used to identify LILR and SPP1 MΦ. Cells were sequentially gated to exclude debris (R1), doublets (R2-R3) and dead cells (R4). CD45+ leukocytes were then gated (R5), followed by exclusion of T, B and plasma cells (R6-R8). CD11b+ myeloid cells were identified (R9) and subdivided into MΦ/Monocyte (CD14+CD163-) and resident MΦ (CD14+CD163+). From the MΦ/Monocyte gate, LILR MΦ were identified as LILRB2+AIF1+ cells based on fluorescence minus one (FMO) controls. From the resident MΦ gate, SPP1 MΦ were identified as SPP1+SDC2+based on FMO controls. B) Flow cytometry gating strategy used for sorting LILR and SPP1 MΦ. Following identification of CD11b+ myeloid cells R9, MΦ/Monocyte and resident MΦ gates were defined as in panel A. LILR MΦ were sorted from the MΦ/Monocyte gate based on LILRB2+ expression as defined by the LILRB2 FMO control. SPP1 MΦ were sorted from the resident MΦ gate based on SDC2+ expression, as defined by the SDC2 FMO control. SDC2- resident MΦ were collected as a comparator population ($SDC2^-$ $LILRB2^-$ MΦ). C) Experimental workflow for functional studies. Single-cell suspensions from 12 CLAD lungs samples (6 BOS, 6 RAS) were thawed stained with sorting flow antibody panel, and subjected to spectral-enhanced fluorescence-activated cell sorting to isolate LILR MΦ, SPP MΦ and $SDC2^-LILRB2^-$ MΦ. Sorted cells were cultured for 24h prior to downstream analyses.

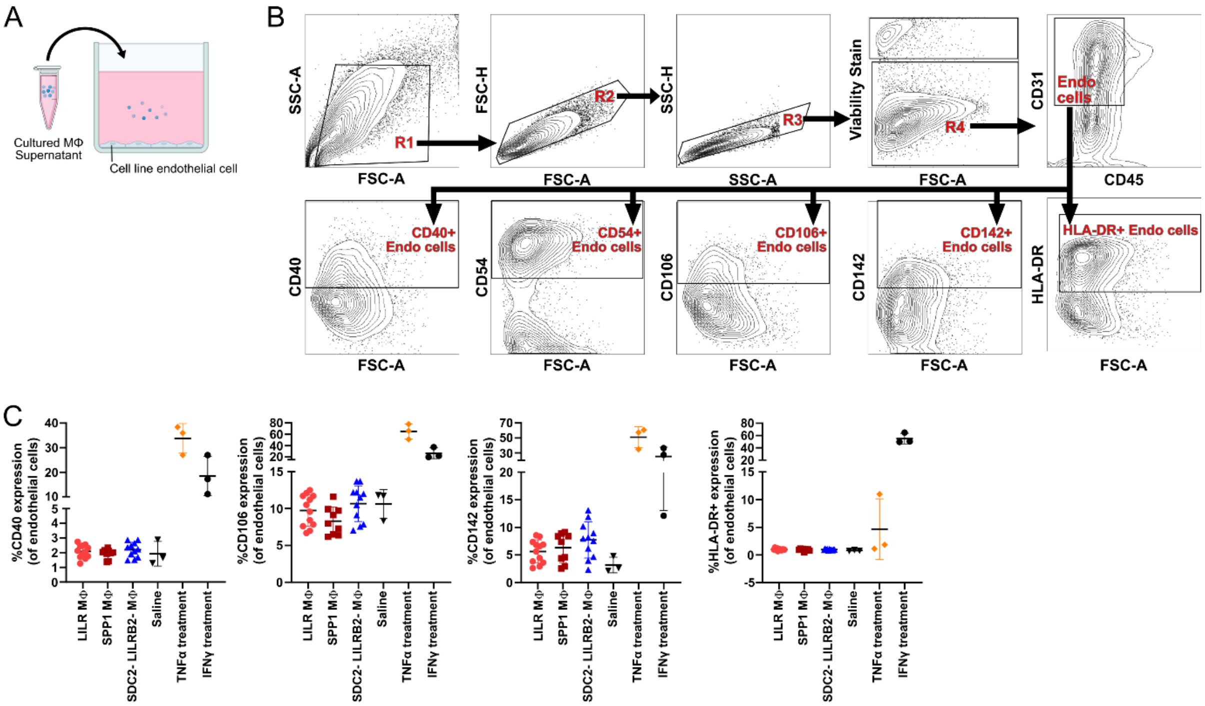


**Supplementary Figure 10: Assessment of endothelial activation following exposure to macrophage-conditioned supernatants.** A) Schematic diagram of in vitro endothelial cell culture following exposure to macrophage-conditioned supernatant obtained from cultured CLAD lung macrophages. B) Flow cytometry gating strategy used to identify endothelial cells and quantify expression of activation markers CD40, CD54, CD106, CD142 and HLA-DR. C) Quantification of endothelial cell activation marker expression across macrophage-conditioned supernatants and control conditions.

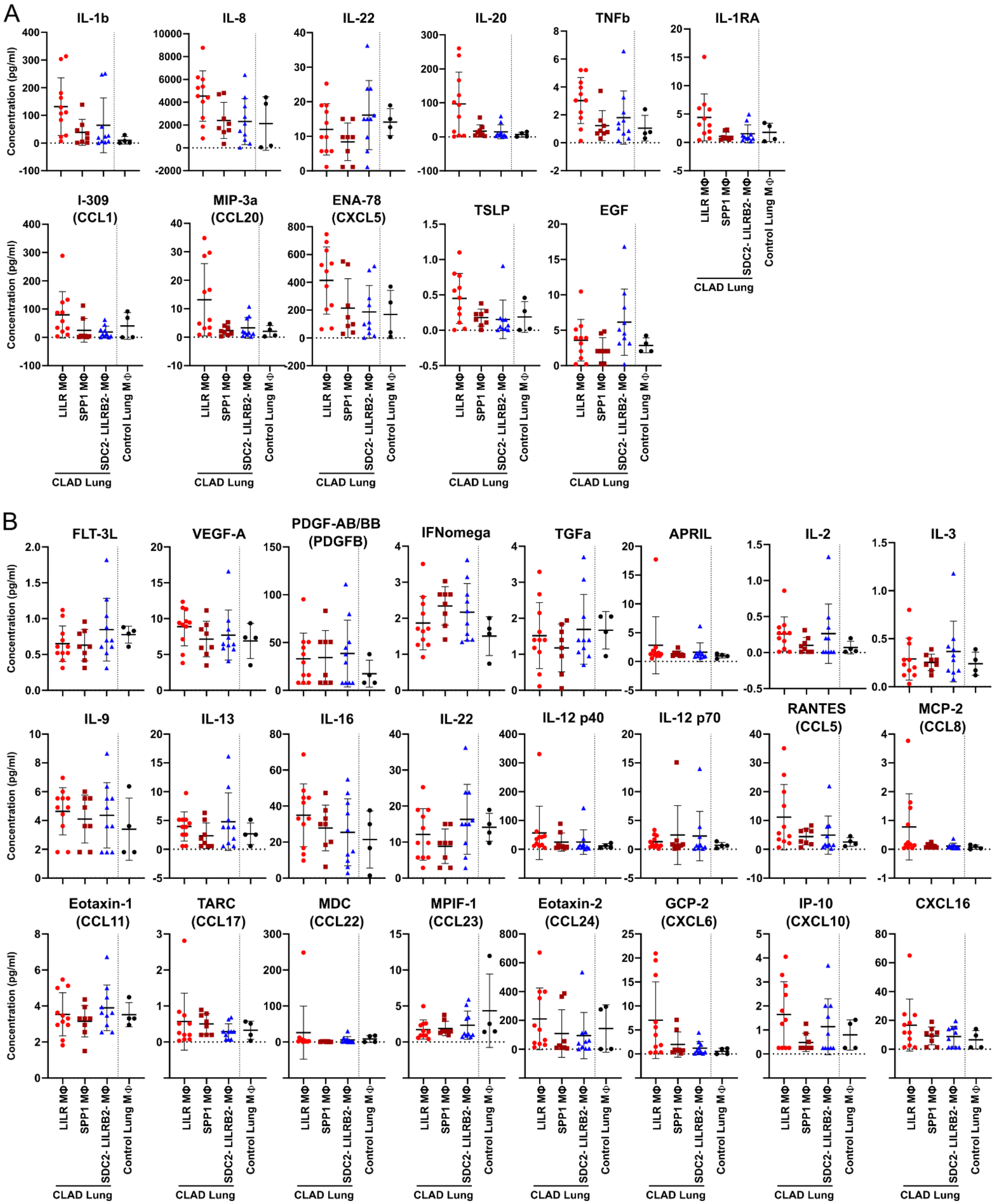
A
IL-1b
IL-8
IL-22
IL-20
TNFb
IL-1RA
I-309 (CCL1)
MIP-3a (CCL20)
ENA-78 (CXCL5)
TSLP
EGF
Concentration (pg/ml)
LILR MΦ
SPP1 MΦ
SDC2- LILRB2- MΦ
Control Lung MΦ
CLAD Lung
B
FLT-3L
VEGF-A
PDGF-AB/BB (PDGFB)
IFNomega
TGFa
APRIL
IL-2
IL-3
IL-9
IL-13
IL-16
IL-22
IL-12 p40
IL-12 p70
RANTES (CCL5)
MCP-2 (CCL8)
Eotaxin-1 (CCL11)
TARC (CCL17)
MDC (CCL22)
MPIF-1 (CCL23)
Eotaxin-2 (CCL24)
GCP-2 (CXCL6)
IP-10 (CXCL10)
CXCL16

**Supplementary Figure 11: Soluble analytes showing trend-level or non-significant differences across macrophage (MΦ) populations.** A) Quantification of multiplex assay analytes from CLAD macrophages culture supernatants demonstrating trend-level differences across LILR, SPP1 and SDC2$^{-}$LILRB2$^{-}$ MΦ populations ($P < 0.1$, $P > 0.05$). B) Quantification of multiplex assay analytes from CLAD macrophage supernatants showing no statistically significant differences across groups ($P > 0.1$). Group comparisons were performed using a Kruskal–Wallis test.

### SUPPLEMENTARY TABLES

#### Table S1 – Genes included in gene module scores

| Module | Genes |
|---|---|
| Pro-inflammatory module | ACOD1, AIF1, AOAH, ARG1, ARID5B, BASP1, CCL3, CCL4, CCRL2, CD274, CIITA, CLEC4E, CTSS, CXCL2, F13A1, FAM126B, FMNL2, FYB1, GADD45A, GBP2, HCAR2, HLA-DRA, HP, IGSF6, IL1B, IL1RN, IL36G, IRGM, LCN2, LILRA5, PLA2G7, PRDX6, RNF213, SASH1, SLFN5 |
| Pro-fibrotic module | ATP6V0D2, CD9, CD63, CHI3L1, CSTB, CTSD, CTSL, CTSS, FABP5, FTH1, GPNMB, LGALS3, MSRB1, PSAP, SGK1, SPP1, TMSB10, TXN1 |
| Engulfment module | ABI3, ARAP2, CDC42, DOCK5, FYN, INPP5D, PIK3CD, PIK3CG, PSTPIP2, RAC1, RHOA, SYK |
| M1-like module | AK3, APOL1, APOL2, APOL3, APOL6, ATF3, BCL2A1, BIRC3, CCL15, CCL19, CCL20, CCL5, CCR7, CHI3L2, CXCL10, CXCL11, CXCL9, EDN1, FAS, GADD45G, HESX1, HSD11B1, IDO1, IGFBP4, IL12B, IL15, IL15RA, IL2RA, IL6, IL7R, INHBA, IRF1, IRF7, NAMPT, OAS2, OASL, PDGFA, PFKFB3, PFKP, PLA1A, PSMB9, PSMA2, PSME2, PTX3, SLC2A6, SLC31A2, SLC7A5, SLCO4A1, SPHK1, TNF, TNFSF10, TYMP, VCAN, XAF1 |
| M2-like module | ADK, ALOX15, CA2, CCL13, CCL18, CCL23, CD209, CD36, CERK, CHN2, CLEC4A, CLEC7A, CTSC, CXCR4, EGR2, FGL2, FN1, GAS7, GPR86, HEXB, HNMT, HRH1, HS3ST1, HS3ST2, IGF1, LIPA, LPAR6, LTA4H, MAF, MRC1, MS4A4A, MS4A6A, MSR1, P2RY14, SEPP1, SLC38A6, SLC4A7, SLCO2B1, TGFBI, TGFBR2, TLR5, TPST2 |

### Table S2 – Flow cytometry antibodies for macrophage panels

| Antibody | Clone | Isotype | Species | Fluorochrome | Dilution | Supplier | Catalog number |
|---|---|---|---|---|---|---|---|
| Validation Sorting Panel | | | | | | | |
| CD11b | M1/70 | IgG2b, κ | Rat | BV510 | 1/20 | BioLegend | 101263 |
| CD45 | HI30 | IgG1, κ | Mouse | BV605 | 1/50 | BD Biosciences | 564047 |
| HLA-DR | L243 | IgG2a, κ | Mouse | APC-Cy7 | 1/100 | BioLegend | 307618 |
| LILRB2 | 287219 | IgG2a | Mouse | BV421 | 1/100 | R&D Systems | FAB2078V |
| SDC2 | 305515 | IgG2B | Rat | APC | 1/100 | R&D Systems | FAB2965A |
| Fixable Viability Stain 700 | n/a | n/a | n/a | n/a | 1/1000 | BD Biosciences | 564997 |
| Analytical Panel | | | | | | | |
| AIF1 | EPR6136(2) | IgG | Rabbit | AF568 | 1/100 | Abcam | Ab221003 |
| CD11b | M1/70 | IgG2b, κ | Rat | BV510 | 1/20 | BioLegend | 101263 |
| CD14 | M5E2 | IgG2a, κ | Mouse | PE-Cy7 | 1/200 | BD Biosciences | 557742 |
| CD163 | GHI/61 | IgG1, κ | Mouse | BUV737 | 1/100 | BD Biosciences | 741863 |
| CD169 | 7-239 | IgG1, κ | Mouse | BV786 | 1/50 | BD Biosciences | 742996 |
| CD19 | HIB19 | IgG1, κ | Mouse | Biotinylated | 1/100 | BioLegend | 302204 |
| CD3 | OKT3 | IgG2a, κ | Mouse | Biotinylated | 1/200 | invitrogen | 13-0037-82 |
| CD45 | HI30 | IgG1, κ | Mouse | BV605 | 1/50 | BD Biosciences | 564047 |
| CD56 | 5.1H11 | IgG1, κ | Mouse | Biotinylated | 1/200 | BioLegend | 362536 |
| CD66b | G10F5 | IgM, κ | Mouse | Biotinylated | 1/100 | BioLegend | 305120 |
| CD71 | CY1G4 | IgG2a, κ | Mouse | PerCP-Cy5.5 | 1/200 | BioLegend | 334114 |
| CXCR4 | 12G5 | IgG2a, κ | Mouse | BUV563 | 1/100 | BD Biosciences | 741400 |
| HLA-DR | L243 | IgG2a, κ | Mouse | APC-Cy7 | 1/100 | BioLegend | 307618 |
| LILRB2 | 287219 | IgG2a | Mouse | BV421 | 1/100 | R&D Systems | FAB2078V |
| LILRB4 | ZM3.8 | IgG1 | Mouse | BUV395 | 1/50 | BD Biosciences | 742811 |
| SDC2 | 305515 | IgG2B | Rat | APC | 1/100 | R&D Systems | FAB2965A |
| SPP1 | 223126 | IgG2A | Mouse | PE | 1/100 | R&D Systems | IC14331P |
| Fixable Viability Stain 700 | n/a | n/a | n/a | n/a | 1/1000 | BD Biosciences | 564997 |
| Culture Sorting Panel | | | | | | | |
| CD11b | M1/70 | IgG2b, κ | Rat | BV510 | 1/20 | BioLegend | 101263 |
| CD138 | MI15 | IgG1, κ | Mouse | AF700 | 1/100 | BioLegend | 356511 |
| CD14 | M5E2 | IgG2a, κ | Mouse | PE-Cy7 | 1/200 | BD Biosciences | 557742 |
| CD163 | GHI/61 | IgG1, κ | Mouse | BUV737 | 1/100 | BD Biosciences | 741863 |
| CD19 | HIB18 | IgG1, κ | Mouse | BV650 | 1/100 | BioLegend | 302238 |
| CD3 | OKT3 | IgG2a, κ | Mouse | PerCP-Cy5.5 | 1/100 | BioLegend | 317336 |
| CD45 | HI30 | IgG1, κ | Mouse | BV605 | 1/50 | BD Biosciences | 564047 |
| EpCAM | KS1/4 | IgG2a | Mouse | BUV563 | 1/50 | BD Biosciences | 748383 |
| HLA-DR | L243 | IgG2a, κ | Mouse | APC-Cy7 | 1/100 | BioLegend | 307618 |
| LILRB2 | 287219 | IgG2a | Mouse | BV421 | 1/100 | R&D Systems | FAB2078V |
| LILRB4 | ZM3.8 | IgG1 | Mouse | BUV395 | 1/50 | BD Biosciences | 742811 |
| SDC2 | 305515 | IgG2B | Rat | APC | 1/100 | R&D Systems | FAB2965A |
| TRAIL | RIK-2 | IgG1, κ | Mouse | PE | 1/100 | BioLegend | 308206 |
| Fixable Viability Stain 620 | n/a | n/a | n/a | n/a | 1/1000 | BD Biosciences | 564996 |

BD Biosciences, San Jose, CA, USA
BioLegend, San Diego, CA, USA
Invitrogen, Waltham, MS, USA
R&D Systems, Minneapolis, MN, USA

### Table S3 – Primers

| Target mRNA | Forward Primer | Reverse Primer |
|---|---|---|
| ACTB | AGCACAGAGCCTCGCCTT | CATCATCCATGGTGAGCTGG |
| AIF1 | CCCTCCAAACTGGAAGGCTTCA | CTTTAGCTCTAGGTGAGTCTTGG |
| LILRB2 | GTGTGGTCTTCACCCAGTGATC | AGCCGACATCAGAGACACACTG |
| MMP9 | GCCACTACTGTGCCTTTGAGTC | CCCTCAGAGAATCGCCAGTACT |
| SDC2 | GCTCCAAAAGTGGAAACCACGAC | ATCCTCTTCGGCTGGGTCCATT |

### Table S4 – Flow cytometry antibodies for endothelial cell phenotypic analysis

| Antibody | Clone | Isotype | Species | Fluorochrome | Dilution | Supplier | Catalog number |
|---|---|---|---|---|---|---|---|
| CD31 | M89D3 | IgG2a | Mouse | BV786 | 1/200 | BD Biosciences | 744467 |
| CD40 | 5C3 | IgG1, κ | Mouse | BV650 | 1/100 | Biolegend | 334338 |
| CD45 | HI30 | IgG1, κ | Mouse | BV510 | 1/100 | BioLegend | 304036 |
| CD54 | HA558 | IgG1, κ | Mouse | PE-Dazzle | 1/100 | Biolegend | 353117 |
| CD106 | STA | IgG1, κ | Mouse | APC | 1/100 | Biolegend | 305810 |
| CD142 | NY2 | IgG1, κ | Mouse | PE | 1/100 | Biolegend | 365204 |
| HLA-DR | L243 | IgG2a, κ | Mouse | APC-Cy7 | 1/100 | Biolegend | 307618 |
| Fixable Viability Stain 620 | n/a | n/a | n/a | n/a | 1/1000 | BD Biosciences | 564996 |

BD Biosciences, San Jose, CA, USA
BioLegend, San Diego, CA, USA
